# CAMSAP2 condensates drive a γ-TuRC-independent pathway for non-centrosomal microtubule nucleation

**DOI:** 10.64898/2026.05.28.728596

**Authors:** Ayhan Yurtsever, Tsuyoshi Imasaki, Ryota Kitano, Kien Xuan Ngo, Tomoaki Yagi, Shuhei Kuno, Hanjin Liu, Takaaki Kato, Hideki Shigematsu, Md Fahim Newaz, Yuji Sugita, Takeshi Fukuma, Ryo Nitta

## Abstract

Microtubule nucleation is commonly viewed as a γ-tubulin ring complex (γ-TuRC)-templated process ^1–3^, yet many differentiated cells build extensive non-centrosomal microtubule arrays of unclear origin ^4–6^. Although spontaneous tubulin nucleation has long been observed in vitro as a γ-TuRC-independent nucleation process ^7–10^, its mechanism and cellular relevance have remained unclear. Here we show that CAMSAP2, a microtubule minus-end-binding protein, links spontaneous nucleation to non-centrosomal microtubule organization. Cryo-electron microscopy (cryo-EM), high-speed atomic force microscopy (hsAFM), and molecular dynamics simulations reveal that CAMSAP2 lowers the nucleation barrier by straightening tubulin oligomers and promoting lateral protofilament interactions that drive sheet formation and closure into microtubules. hsAFM captures stepwise nucleation and early growth, revealing that tubulin rings can serve as productive intermediates rather than dead-end depolymerization products. CAMSAP2 further self-organizes through liquid–liquid phase separation (LLPS) to concentrate tubulin and assembly intermediates, thereby promoting efficient microtubule formation at non-centrosomal microtubule-organizing centers. In HeLa cells, dispersed CAMSAP2 condensates act as γ-TuRC-independent microtubule-organizing centers alongside centrosomal asters, supporting a model in which spontaneous nucleation contributes to non-centrosomal microtubule biogenesis.

## INTRODUCTION

Microtubules are polarized cytoskeletal polymers built from αβ-tubulin heterodimers. Their plus ends are highly dynamic, whereas minus ends are often capped or stabilized by microtubule-associated proteins (MAPs) ^11–15^. Microtubule nucleation, the initial formation of a polymer from tubulin dimers, faces a substantial kinetic barrier in cells ^16^. In contrast to actin, microtubules must establish longitudinal contacts along protofilaments, lateral contacts between neighboring protofilaments, and finally undergo tube closure ^7^. Early in vitro studies on purified tubulin showed that microtubules can nucleate spontaneously at high tubulin concentrations and that tubulin first forms rings and short, curved oligomers (**Supplementary Fig. 1a)**. More recent structural and mutational studies have reinforced the view that the abundance and straightness of these intermediates are critical determinants of nucleation efficiency ^8–10,17–19^. Consistent with this idea, recent in situ cryo-electron tomography (cryo-ET) of regenerating axons revealed abundant clusters of curled protofilaments at elongating microtubule tips, suggesting that structural remodeling of tubulin oligomers is a critical but poorly understood step in nucleation ^20^, yet how these heterogeneous oligomers are structurally reorganized into a productive nucleus and how their local conformational states influence early microtubule growth remain unclear.

In cells, microtubule nucleation is most commonly attributed to the γ-TuRC, which provides a structural template that promotes nucleation and stabilizes the minus end ^1–3^. However, many differentiated cells, including neurons, epithelial cells, and muscle cells, assemble extensive γ-TuRC independent non-centrosomal microtubule arrays that persist even when γ-TuRC activity is spatially restricted or reduced ^4–6^. These observations raise the possibility that microtubules can be generated through γ-TuRC–independent mechanisms.

Several MAPs, including XMAP215/chTOG, TPX2, CLASP, p150Glued, doublecortin, and GAS2 family proteins, have been reported to promote microtubule assembly by accelerating polymerization, stabilizing early intermediates, or organizing protofilament interactions ^4,6,21–27^. Yet, these activities are generally thought to facilitate growth from pre-existing or transient nuclei rather than to generate stable, elongation-competent microtubule seeds independently of γ-tubulin. Thus, despite extensive studies on MAP-mediated regulation of microtubule dynamics, the molecular mechanism by which cells overcome the nucleation barrier in the absence of γ-TuRC remains unresolved. How these heterogeneous, non–γ-TuRC-templated intermediates are converted into productive nuclei in cells remains unknown.

CAMSAP/Patronin proteins are minus-end binding MAPs that are central to the organization of non-centrosomal microtubule arrays in animals ^6,28–31^. They recognize and protect free minus ends, suppress minus-end dynamics, and help build non-centrosomal microtubule networks in epithelial cells, neurons, and other polarized cell types. We previously showed that CAMSAP2 undergoes LLPS, robustly recruits tubulin, and behaves as a γ-TuRC–independent microtubule-organizing center that nucleates microtubules and produces radial Cam2-asters in vitro ^32^. Recent work has also highlighted a post-nucleation role of CAMSAP family proteins: Rai et al. showed that CAMSAP2/3 can bind newly formed minus ends of γ-TuRC-nucleated microtubules and promote their release from γ-TuRC, thereby acting as organizers of free non-centrosomal minus ends ^33^. However, this interpretation does not exclude a role for CAMSAPs in nucleation itself. Rather, their ability to engage newly formed minus ends and promote elongation and stabilization suggests that CAMSAPs may support a continuous pathway spanning nucleation and immediate post-nucleation maturation. These condensates likely mimic non-centrosomal microtubule organizing centers (ncMTOCs) in cells, where CAMSAP2 or CAMSAP3 concentrate tubulin and minus ends at the Golgi, cortex, or apical membrane ^30,34,35^.

Structurally, CAMSAP family proteins contain N-terminal regions that mediate self-association, LLPS, and tubulin binding, and a C-terminal CKK domain that binds the microtubule lattice and selectively recognizes minus ends ^36,37^. The CKK domain loop 7 inserts into the inter-protofilament groove and contacts the tubulin H11–H12 region, suggesting a potential role in stabilizing lateral interactions within specific lattice geometries. Despite these advances, it remains unclear how CAMSAPs remodel heterogeneous tubulin oligomers into nucleation-competent seeds, whether CAMSAP-driven assemblies represent a bona fide γ-TuRC–independent nucleation pathway, and how such activity contributes to non-centrosomal microtubule network formation in cells. To address these questions, we combined in vitro reconstitution with cryo-EM, hsAFM, molecular dynamics (MD) simulations, and targeted mutagenesis to define how CAMSAP2 reshapes tubulin oligomers to drive spontaneous nucleation and early minus-end elongation. We then examined how these molecular events give rise to higher-order aster-like assemblies and, finally, established a live-cell system to directly compare γ-TuRC–dependent and γ-TuRC–independent microtubule network assembly, demonstrating that CAMSAP2 condensates function as ncMTOCs in living cells.

## RESULTS

### Cryo-EM reveals spontaneous microtubule nucleation in CAMSAP2-LLPS

We previously showed that CAMSAP2 nucleates microtubules via liquid–liquid phase separation ^32^. To determine whether CAMSAP2 merely concentrates tubulin or actively organizes specific nucleation intermediates, we re-examined early assembly events by cryo-EM using the CAMSAP2 CC1–CKK fragment (hereafter CC1-CKK), which shows less condensed nucleation center than the full-length protein ^32^ and thus permits capturing the states cryo-EM. Tubulin and CC1–CKK (10:1 molar ratio; 30 µM tubulin, 3 µM CC1–CKK) were co-incubated on ice for 30 min, shifted to 37 °C for 30 s or 1 min, and plunge-frozen. Cryo-electron tomography (ET) datasets denoised using Cryo-CARE ^38–40^ revealed short curved protofilaments, rings, sheets, and elongating microtubules with curled ends and sheet–ring junctions (**Fig. 1a,b, Supplementary Fig. 1**). At 1 min, we also observed Cam2-aster formation, consistent with our previous in vitro reconstitution ^32^.

**Fig. 1.**
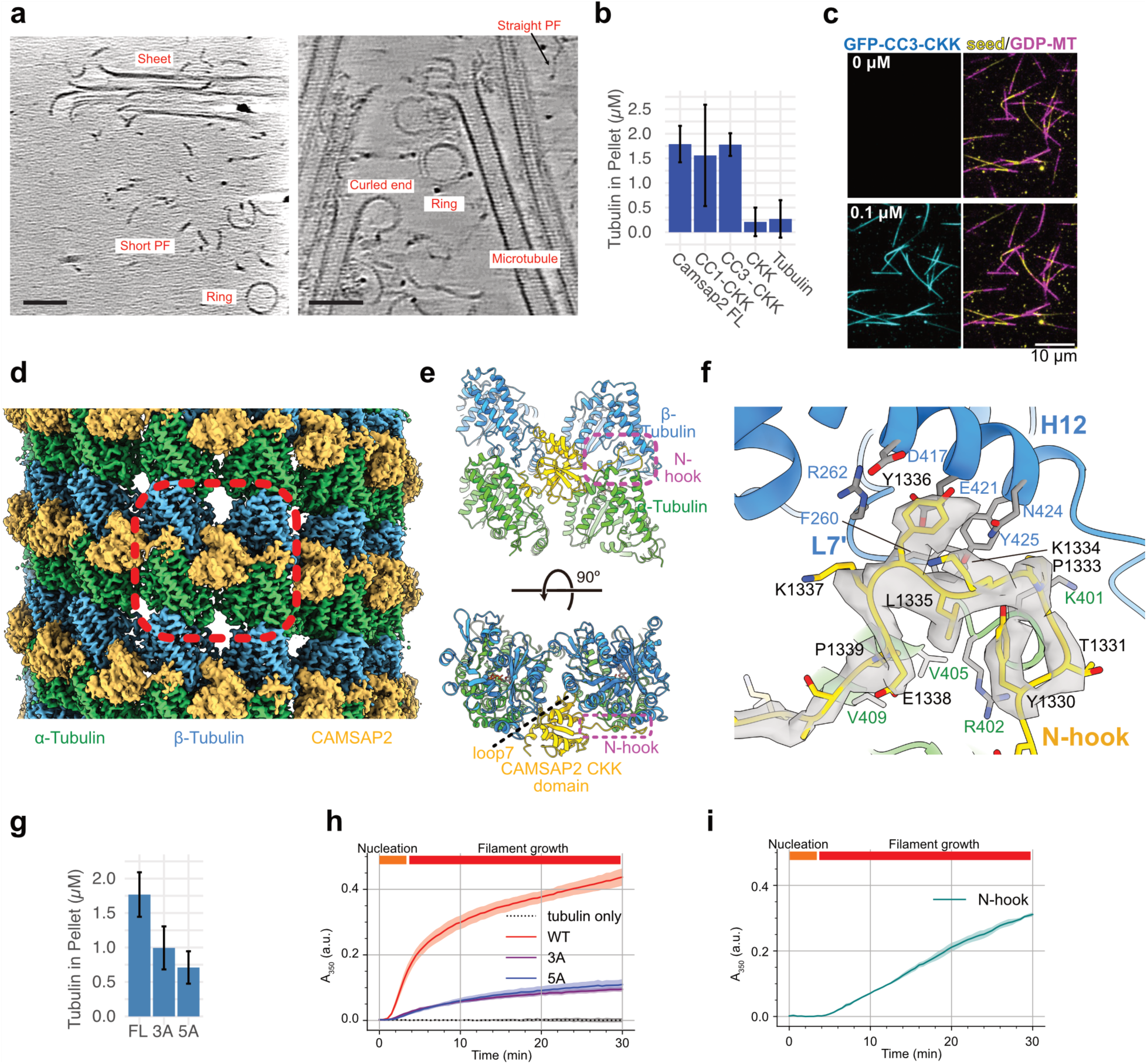
Cryo-EM structure of CAMSAP2-microtubule complex. **a**, Cryo-ET reconstructions of 30 µM tubulin with 3 µM CAMSAP2 CC1–CKK after 30 s (left) or 1 min (right), denoised with cryo-CARE. Representative structures include tubulin sheets, rings, short protofilaments, and microtubules with curled tips. Scale bars, 50 nm. **b**, Microtubule stabilization assay. **c**. TIRF observation of the microtubules before and after perfusing CC3-CKK in the PEM buffer. GFP-CC3-CKK is in cyan, microtubule seed in yellow, GDP-microtubule in magenta. **d**. Overall structure of the microtubule with CAMSAP2 CC3-CKK. α-Tubulin is in green, β-Tubulin in cyan, and CAMSAP2 in yellow. The red dashed outline indicates the region modeled in (e). **e**. Cartoon representation of the two tubulin heterodimers laterally connected by CAMSAP2 CKK domain and N-hook. The purple dashed region marks the N-hook and corresponds to the region shown in (f). **f**. Molecular interaction between N-hook and tubulin heterodimer. The density of the N-hook is shown in gray. **g**. Stabilization assay with CAMSAP2 FL or with CAMSAP2 FL N-hook mutants. **h**. Turbidity assay of 8.5 µM tubulin in the buffer PEM-GTP, monitoring optical density (OD) at 350 nm over 30 minutes to assess microtubule polymerization. The experiment was performed under four conditions: (1) tubulin alone (without CAMSAP2), (2) tubulin with 0.5 µM wild-type CAMSAP2 GFP-CC1-CKK, (3) tubulin with 0.5 µM CAMSAP2 GFP-CC1-CKK carrying the N-hook 3A mutation, and (4) 0.5 µM CAMSAP2 GFP-CC1-CKK carrying the 5A mutation. Each experiment was repeated three times. I. Tubulin turbidity assay of 8.5 µM of tubulin with 2 µM of GFP-CC3-N-hook.

These intermediates were observed at very early time points under conditions where tubulin alone does not nucleate, indicating that the observed rings and curved oligomers are unlikely to represent depolymerization products. Instead, their structural continuity with elongating microtubules suggests that they function as assembly intermediates. We further observed ring-to-higher-order transitions, including stacked and partially opened rings and their conversion into sheet-like assemblies (**Fig. 1a, Supplementary Fig. 1b-d**). Similar incorporation of oligomeric intermediates into growing lattices has been described in other MAP-assisted nucleation systems, supporting a model in which such structures directly contribute to microtubule assembly ^10,41,42^.

These observations are consistent with a spontaneous nucleation mechanism in which tubulin assembly proceeds through two coupled steps: formation of a critical nucleus from short longitudinal oligomers and subsequent filament growth through sheet elongation and tube closure (**Supplementary Fig. 1a: Nucleation & Filament growth**)^8–10,17,42–44^. In this framework, CAMSAP2 does not template nucleation in a γ-TuRC-like manner; instead, it organizes and remodels intrinsic tubulin oligomers to promote their transition into elongation-competent intermediates.

### N-hook and CKK domains reinforce longitudinal and lateral protofilament contacts

To investigate how CAMSAP2 recognizes tubulin assemblies at atomic resolution, we determined the structure of the CAMSAP-microtubule complex by cryo-EM single particle analysis (SPA) using a CC3-CKK construct containing the CC3 coiled-coil and CKK domains (**Supplementary Fig. 2, S3a**). This construct retains robust microtubule nucleation and stabilization activity while exhibiting reduced minus-end bias (**Fig. 1b,c, Supplementary Fig. 3b,c**) ^32^, enabling dense decoration of CAMSAP2 suitable for high-resolution reconstruction.

The CKK domain binds at the intra-dimer interface of tubulin subunits and mediates lateral contacts between adjacent protofilaments, except at the microtubule seam (**Fig. 1d,e, Supplementary Fig. 3d**) ^36,37^. By wedging between neighboring protofilaments, the CKK domain stabilizes the tubular lattice and contributes to defining microtubule architecture (**Fig. 1e, bottom**). Structural comparison confirmed that the CAMSAP2 CKK domain closely resembles those of CAMSAP1 and CAMSAP3 (RMSD 1.191 and 1.031 against PDB 6QVJ and 5M5C), highlighting a conserved binding mode across the CAMSAP family.

Although the CC3 domain itself was not resolved due to its flexibility, we identified a prominent hook-like density extending from the CKK domain, corresponding to the conserved linker between CC3 and CKK (**Fig. 1d-f, Supplementary Fig. 3e**). This element, which we term the “N-hook,” adopts a curved conformation that inserts into the intra-dimer interface of the tubulin heterodimer. The N-hook engages the groove between α- and β-tubulins and is stabilized by conserved residues (Y1330, P1333, L1335, and P1339), thereby reinforcing longitudinal contacts within PFs (**Fig. 1f, Supplementary Fig. 3e**).

Together with the lateral stabilization mediated by the CKK-domain, the N-hook provides a complementary mechanism that strengthens protofilament integrity along both longitudinal and lateral axes. These findings reveal a dual structural basis by which CAMSAP stabilizes early assembly intermediates and promotes their progression toward elongation-competent microtubules (**Fig. 1d**).

### N-hook drives microtubule nucleation and filament growth

To assess the functional contribution of the N-hook, we generated two mutants in the CC1–CKK context (CAMSAP2-CC1–CKK-3A and -5A), in which conserved N-hook residues were replaced, and examined their effects on microtubule nucleation and stabilization (**Supplementary Fig. 3a,f**).

In a microtubule stabilization assay, both mutants showed reduced activity compared to wild type, with ∼40% and ∼60% reductions for the 3A and 5A mutants, respectively, indicating that the N-hook makes an essential contribution to lattice stability (**Fig. 1g**). We next assessed nucleation by turbidity at 350 nm ^45^. Both mutants displayed a prolonged lag phase, reduced initial polymerization rates, and decreased secondary growth, whereas tubulin alone did not polymerize detectably (**Fig. 1h**).

Consistent with these results, negative-stain EM revealed that under near-physiological ionic strength (PEM + 50 mM KCl), the 3A mutant generated smaller asters, whereas the 5A mutant was severely impaired, yielding only sparse aster-like structures (**Supplementary Fig. 4a-c**). These data show that the N-hook becomes increasingly critical for efficient nucleation and growth under physiological conditions

To isolate the N-hook contribution from that of the CKK, we analyzed an N-hook–only construct, GFP–CC3–N-hook. Turbidity assays below the critical concentration revealed that GFP–CC3–N-hook retains intrinsic polymerization activity: although the rapid initial phase observed with wild-type CC1–CKK was diminished, delayed but sustained microtubule assembly emerged after several minutes (**Fig. 1h,i**). The subsequent polymerization slope was comparable to that of wild-type CC1–CKK and exceeded that of the N-hook mutants, indicating that the N-hook alone confers weak but intrinsic nucleation competence. Negative-stain EM confirmed that GFP–CC3–N-hook promotes microtubule formation from small condensates but rarely produces Cam2-aster–like structures (**Supplementary Fig. 4d**). We also found that CC3–CKK and CKK exhibit lower phase-separation propensity than the others (**Supplementary Fig. 4e-i**).

This division of labor resembles previously described synergistic nucleation systems ^27^. Taken together, these results establish that the N-hook is indispensable for efficient microtubule nucleation and growth. By reinforcing longitudinal protofilament contacts and cooperating with CKK-mediated lateral stabilization, the N-hook enables CAMSAP2 to reach full nucleation efficiency and sustain filament growth.

### Molecular visualization of early nucleation reveals N-hook-driven ring-to-protofilament conversion

We used hsAFM to visualize how CAMSAP2 promotes microtubule nucleation at molecular resolution. Application of 1 µM tubulin to the mica produced numerous rings structures (**Fig. 2a, Supplementary Video 1, Supplementary Fig. 5a**). These rings, composed of longitudinally aligned tubulin dimers, were stabilized by the mica surface and remained intact for >10 min at room temperature, limiting further polymerization ^46^.

**Fig. 2.**
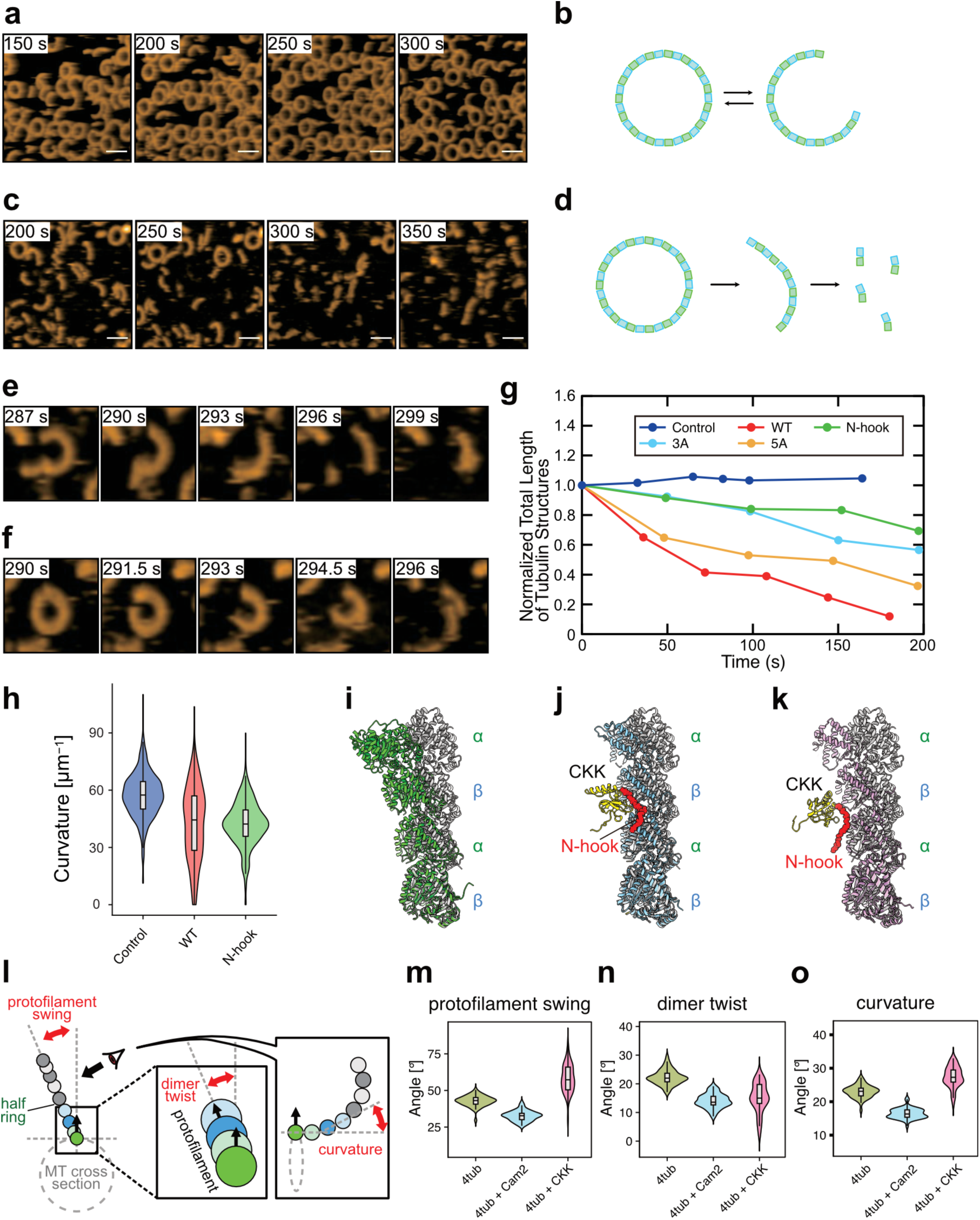
hsAFM and MD simulations reveal CAMSAP2-driven straightening of tubulin oligomers. hsAFM movies showing dynamic changes in tubulin ring curvature in the (a) absence and (c) presence of CAMSAP2 CC1–CKK. Tubulin and CAMSAP2 CC1–CKK or its mutants were used at 1 µM and 0.1 µM, respectively (see also **Supplementary Video 1 and 2**). **a**, Tubulin rings on the mica surface imaged by hsAFM. **b**, Schematic representation of tubulin ring on a mica surface. **c**, Tubulin rings captured after addition of CAMSAP2. **d**, Schematic representation of tubulin ring with CAMSAP2 on a mica surface. **e, f**, Representative movie frames showing the disruption of rings and the straightening and shortening of short protofilaments following CAMSAP2 addition. **g**, Ring disassembly rate quantified by the normalized total length of tubulin structures: control (tubulin only, blue), WT (red), N-hook (green), 3A (cyan), and 5A(yellow). See **Supplementary** Fig. 5e for more details. **h**, Box plot of tubulin oligomer (length < 50 nm) curvature under identical states: control (tubulin only, blue), WT (red), and N-hook (green). Mean ± SD = 57.18 ± 13.82 μm⁻¹ (N=463) for control; 40.44 ± 20.39 μm⁻¹ (N=368) for WT; 42.66 ± 15.41 μm⁻¹ (N=186) for N-hook. Two-sample t-tests confirm significant differences between control and WT (p ≤ 1.26 × 10⁻⁴⁰) and between control and N-hook (p ≤ 4.27 × 10⁻³⁰) at p ≤ 0.05, but no significant difference between WT and N-hook (p ≤ 0.19). See **Supplementary** Fig. 5f for more details. Scale bars, 50 nm. Imaging rates: (a) 1 s/frame; (b–d) 1.5 s/frame. **i-k**, Representative snapshots from all-atom MD simulations of (i) a bare tubulin tetramer (green), (j) a tubulin tetramer (cyan) bound by CAMSAP2 N-hook(red)–CKK (yellow), and (k) a tubulin tetramer (pink) in which the CAMSAP2 N-hook (red) has dissociated from CKK (yellow), each superimposed on a straight protofilament (gray). Snapshots are shown after 1.0 µs of simulation. Three independent simulations were performed for the bare tetramer and the N-hook–CKK–bound tetramer, and one simulation for the N-hook–dissociated state. **l**, Schematic illustration of the geometric parameters used to describe tubulin tetramers. The arrow originating from the tubulin (circle) represents the vector that is perpendicular to the microtubule surface. **m-o**, Violin plots with overlaid box plots showing the distributions of the parameters defined in (l), calculated from 50 MD samples per state: (m) protofilament swing, (n) dimer twist, and (o) curvature.

Addition of 0.1 µM CC1–CKK rendered these rings highly dynamic: they fragmented into semicircular structures, transitioned into gently curved single protofilaments, and eventually disassembled (**Fig. 2c,d,f, Supplementary Video 1**). To assess the contribution of the N-hook, we examined N-hook mutants (3A, 5A) with reduced nucleation activity (**Supplementary Fig. 5b,c**). Both mutants showed slower ring disassembly than wild type, implicating that the N-hook promotes curvature reduction and ring-to-protofilament conversion (**Supplementary Fig. 5b,c, Supplementary Video 2**). Consistent with this, an isolated N-hook construct (CC3–N-hook or N-hook), lacking CC1 and CKK, was sufficient to induce ring disruption, albeit less efficiently than wild-type CC1–CKK (**Fig. 2g, Supplementary Fig. 5d, Supplementary Video 2**).

Quantitative analysis of curvature and contour-length distributions revealed that both wild-type and the N-hook significantly reduced protofilament curvature to similar extents, accompanied by a decrease in stable rings and an increase in straightened protofilaments (**Fig. 2h, Supplementary Fig. 5e-j**). The N-hook alone produced longer, more persistent straight protofilaments, whereas such species were less abundant and more transient in the presence of wild-type CC1–CKK, suggesting additional regulation by the CKK domain.

Together, these results demonstrate that the N-hook drives the conversion of stable tubulin rings into straightened protofilaments, a key intermediate step toward nucleation. This curvature reduction provides a mechanistic basis for how CAMSAP2 lowers the nucleation barrier and promotes spontaneous microtubule assembly.

### Molecular dynamics reveals N-hook-driven protofilament straightening

To test whether the CAMSAP2 N-hook contributes to PF straightening, we performed all-atom molecular dynamics (MD) simulations on a GDP-state tubulin tetramer representing a short PF segment (**Supplementary Fig. 6**). Simulations were carried out with the GENESIS platform ^47–49^. Two initial models were built from the cryo-EM structure of CAMSAP2 bound to microtubules: (i) a bare tetramer (4tub), composed of a central α, β-heterodimer flanked by α-tubulin (plus end) and β-tubulin (minus end), and (ii) CAMSAP2-bound tetramer (4tub–Cam2), in which the N-hook–CKK module engages the central heterodimer (**Fig. 2i,j**). In one 4tub–Cam2 trajectory, the N-hook spontaneously disengaged from the tubulin groove; this event was treated as a separate state (iii) (**Fig. 2k**).

PF geometry was quantified using three parameters: swing (the angle between the PF axis and the microtubule axis), twist (the rotation between adjacent tubulin dimers within a PF), and curvature (the degree of PF peeling from the microtubule lattice) (**Fig. 2l, Supplementary Fig. 7a,b**). In the absence of stable N-hook engagement (trajectories (i) and (iii)), PFs adopted conformations characteristic of depolymerizing states, with large swing and curvature values (swing: 4tub, 42.9 ± 3.59°; 4tub + CKK, 58.2 ± 9.48°; curvature: 4tub, 23.1 ± 1.86°; 4tub +CKK, 27.4 ± 3.12°) (**Fig. 2m-o, Supplementary Fig. 7**) ^41^. By contrast, the N-hook–bound state (ii) exhibited significantly reduced swing and curvature (swing: 4tub +Cam2, 32.7 ± 3.51°; curvature: 4tub +Cam2, 16.5 ± 1.51°), consistent with N-hook–dependent stabilization of a straighter, microtubule-like PF geometry.

Twist was reduced in all CAMSAP2-containing simulations (ii and iii), including the N-hook-disengaged trajectory, suggesting that the CKK domain primarily suppresses PF twist (**Fig. 2m-o, Supplementary Fig. 7**). By contrast, reduction of swing and curvature depended on stable N-hook engagement, identifying the N-hook as the principal determinant of PF straightening.

Notably, the N-hook–CKK module remained stably associated with a single PF throughout the simulations, supporting the idea that CAMSAP2 can directly act on ring-like or single-PF intermediates. Together, these simulations indicate that CAMSAP2 promotes PF straightening through a division of labor: the N-hook reduces PF curvature and swing, while the CKK domain constrains twist. This coordinated remodeling converts curved oligomers into elongation-competent intermediates, providing a structural basis for CAMSAP2-driven spontaneous microtubule nucleation.

### CAMSAP2 promotes lateral protofilament contacts and sheet closure during filament growth

To visualize CAMSAP2-dependent filament growth by hsAFM, we replaced the mica with a lipid membrane, because the strong negative charge of mica stabilizes tubulin rings but suppresses lateral PF contacts. A lipid mixture of DPPC: DPTAP (92.5:7.5 (wt/wt)) was identified as optimal.

We first examined 20 µM tubulin alone. hsAFM revealed abundant tubulin rings and isolated, straight PFs; although some PFs occasionally approached each other, they failed to establish stable lateral contacts (**Supplementary Fig. 8a, Supplementary Video 3**). All filaments showed a uniform height of ∼4 nm, consistent with single PFs (**Supplementary Fig. 8b**). Thus, tubulin alone on the lipid surface does not progress beyond ring and PF intermediates, likely because lateral association requires partial detachment from the membrane, which is energetically unfavorable.

Upon addition of 2 µM CC1–CKK, polymerization was rapidly initiated, yielding PFs, open tubulin sheets, and microtubules across the field (**Supplementary Fig. 8c, Supplementary Video 3**). Because assembly proceeded quickly, the very earliest transitions were rarely captured; however, intermediate structures were frequently observed. Height measurements confirmed the presence of microtubules (22-24 nm) in regions where filaments were not stacked (**Supplementary Fig. 8d**). As assembly progressed, dense microtubule networks formed, reducing the number of observable intermediates, consistent with rapid consumption of tubulin into higher-order structures.

In areas where assembly was still ongoing, we directly visualized sheet-to-microtubule transitions (**Fig. 3a, Supplementary Video 4**). Elongating sheets displayed heights of ∼15–17 nm (**Fig. 3b**), consistent with open sheet structures, and were surrounded by numerous tubulin rings (Supplementary Fig. 8). Curved PFs were recruited to sheet edges, where they fluctuated between curved and straight conformations before straightening and docking into the lattice (**Fig. 3a**, arrows). This incorporation coincided with measurable increases in sheet width (T-U vs V-W; **Fig. 3a,b**). Once a sheet reached a critical width, it abruptly twisted and closed into a microtubule (**Fig. 3a**), as indicated by a height increase to 22-24 nm (**Fig. 3c**). This sheet-to-tube transition occurred too quickly to be fully resolved by hsAFM. After closure, the proximal region adopted a microtubule-like height, while the distal edge remained slightly lower, consistent with continued sheet-like elongation (1577 s; **Fig. 3a**, yellow). Throughout these processes, tubulin rings and free PFs remained abundant in the background, suggesting that they constitute a local reservoir of subunits for further elongation (**Supplementary Fig. 8e**).

**Fig. 3.**
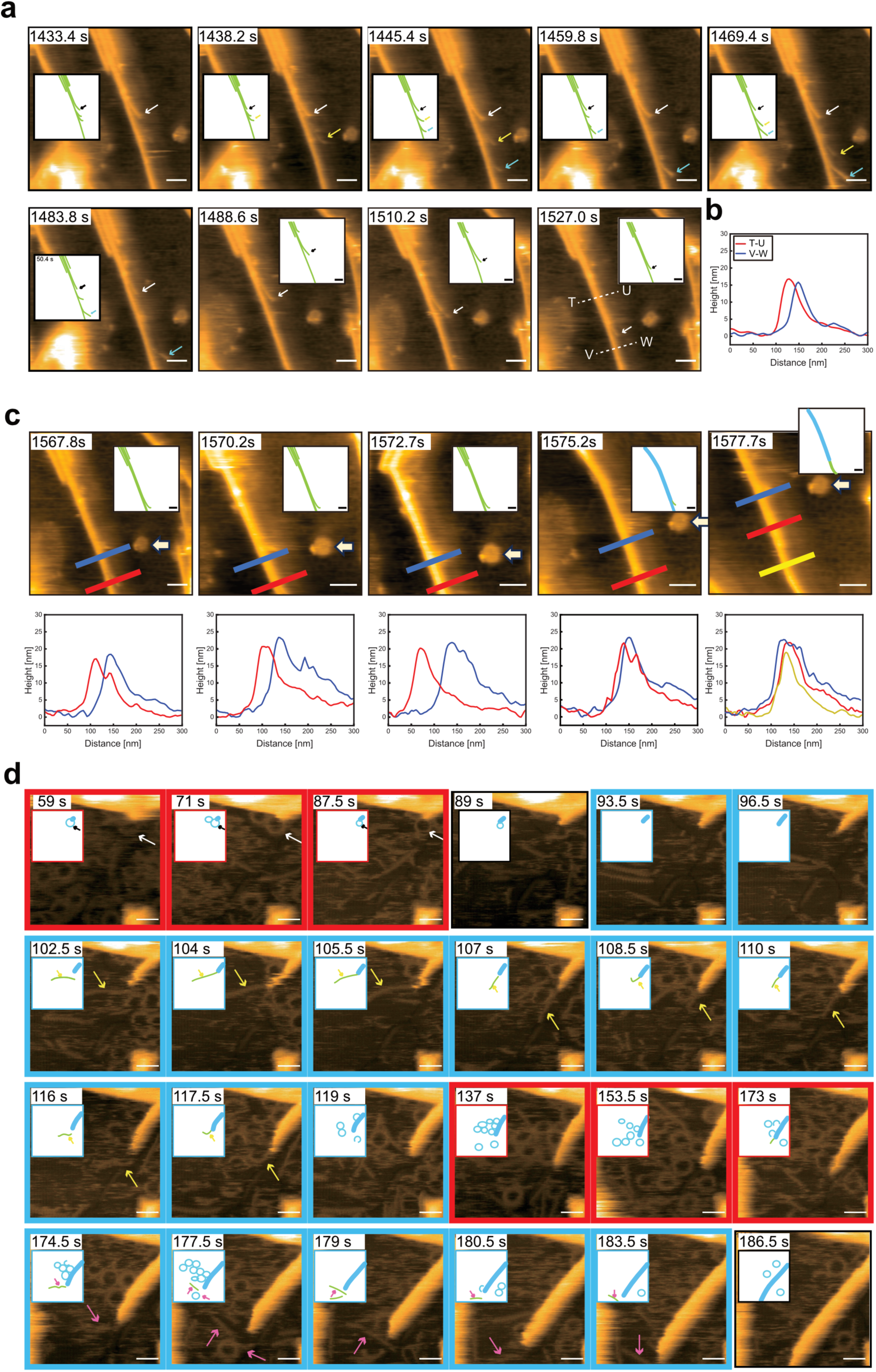
hsAFM reveals CAMSAP2-driven stepwise microtubule assembly on lipid membranes. Representative hsAFM images showing intermediate tubulin structures, protofilament incorporation into sheets, and sheet-to-microtubule transitions on a lipid membrane. Tubulin and CAMSAP2 CC1–CKK were imaged by hsAFM. Protofilaments are color-coded to facilitate tracking across frames; the same protofilament is shown in the same color in all frames (green, light green, and dark green). Rings are colored cyan. **a.** hsAFM images showing gradual incorporation of protofilaments into tubulin sheets through inchworm-like straightening and winding (see also **Supplementary Video 4**). Tubulin and CAMSAP2 CC1–CKK concentrations were 20 µM and 2 µM, respectively. **b.** Cross-sectional profiles of the heights and widths of the tubulin sheets composed of several protofilaments in (A 1527.0 s) T-U and V-W. **c.** Sheet-to-microtubule conversion and cross-sectional profiles of the corresponding heights and widths (see also **Supplementary Video 4**). Arrows indicate fiducial markers, denoting the same position across frames. **d.** Microtubule tips incubated with the CAMSAP2 CC1–CKK 5A mutant. Cyan boxes indicate elongation phases, whereas red boxes indicate pause phases with alternating growth and shrinkage. Arrows indicate tubulin rings or ring stacks engaging the microtubule tip. Tubulin and the 5A mutant were used at 20 µM and 2 µM, respectively (see also **Supplementary Video 5**). Scale bar, 50 nm. Imaging speeds: (a) 5 s/frame, (c) 2 s/frame, and (d) 1.5 s/frame.

Together, these observations show that CAMSAP2 promotes microtubule formation not only by initiating nucleation but also by actively enhancing lateral PF contacts and sheet closure. By facilitating the transition from isolated PFs to laterally associated sheets that rapidly close into tubes, CAMSAP2 bridges nucleation and filament growth, enabling robust microtubule assembly under near-physiological conditions.

### N-hook mutant reveals stepwise microtubule assembly from tubulin rings

Because N-hook mutations impair nucleation, PF straightening, and polymerization (**Fig. 1,2**), we hypothesized that they would attenuate CAMSAP2-driven assembly, thereby enabling visualization of transient intermediates. We therefore performed hsAFM imaging using CAMSAP2 CC1–CKK with 3A or 5A mutations.

Consistent with their reduced activity, neither mutant produced discernible Cam2-Asters even after prolonged incubation. This agrees with negative-stain EM and turbidity assays under near-physiological conditions (PEM + 50 mM KCl), which showed reduced nucleation (**Supplementary Fig. 4a-c**) and slower polymerization (**Fig. 1h**). The slowed kinetics allowed direct visualization of intermediate states during microtubule assembly.

hsAFM revealed that microtubule growth proceeds through stepwise incorporation of tubulin rings and protofilaments into (**Fig. 3d, Supplementary Video 5**). Tubulin rings were frequently aligned along elongating microtubules and then disappeared as the lattice advanced, consistent with incorporation into the growing structure. A similar ring alignment was also observed by cryo-ET at comparable stages (**Fig. 1a, Supplementary Fig. 1b**).

To quantify these dynamics, we segmented trajectories into elongation (cyan) and pause (red) phases (**Fig. 3d**). In a representative movie, 0–30 s corresponded to a pause, 30–60 s to elongation, 60–115 s to a pause, followed by renewed elongation. Elongation phases correlated with the presence of straight or weakly curved oligomeric fragments at a microtubule growth end, whereas pauses coincided with ring-like structures contacting the tip. These observations suggest that the structural state of tubulin assemblies––ring-like versus straightened PF––determines their probability of incorporation ^50^.

Notably, microtubule elongation occurred in discrete steps whose size approximates the circumference of a single tubulin ring (∼250 nm, corresponding to ∼26-30 tubulin subunits) ^32^. Together, these data provide direct dynamic evidence that tubulin rings are not dead-end intermediates but are instead modular building blocks that are incorporated stepwise into growing microtubules.

### CAMSAP2 condensates drive aster network formation in vitro

We next asked how CAMSAP2 assemblies organize higher-order microtubule networks. Because CAMSAP2 forms liquid-like condensates with tubulin ^32^, we used hsAFM on lipid membranes to visualize Cam2-aster formation in real time.

Microtubules radiated from condensates and exhibited dynamic behavior, frequently switching between growth and shrinkage (**Fig. 4a,b, Supplementary Fig. 10, Supplementary Video 6**). When neighboring asters approached, they often fused into a single larger structure (**Fig. 4a**), consistent with condensate fusion observed previously in vitro and in cells (**Supplementary Fig. 10b**) ^32^. In other cases, asters remained distinct but became connected by microtubule bundles, forming an interconnected network (**Fig. 4b**).

**Fig. 4.**
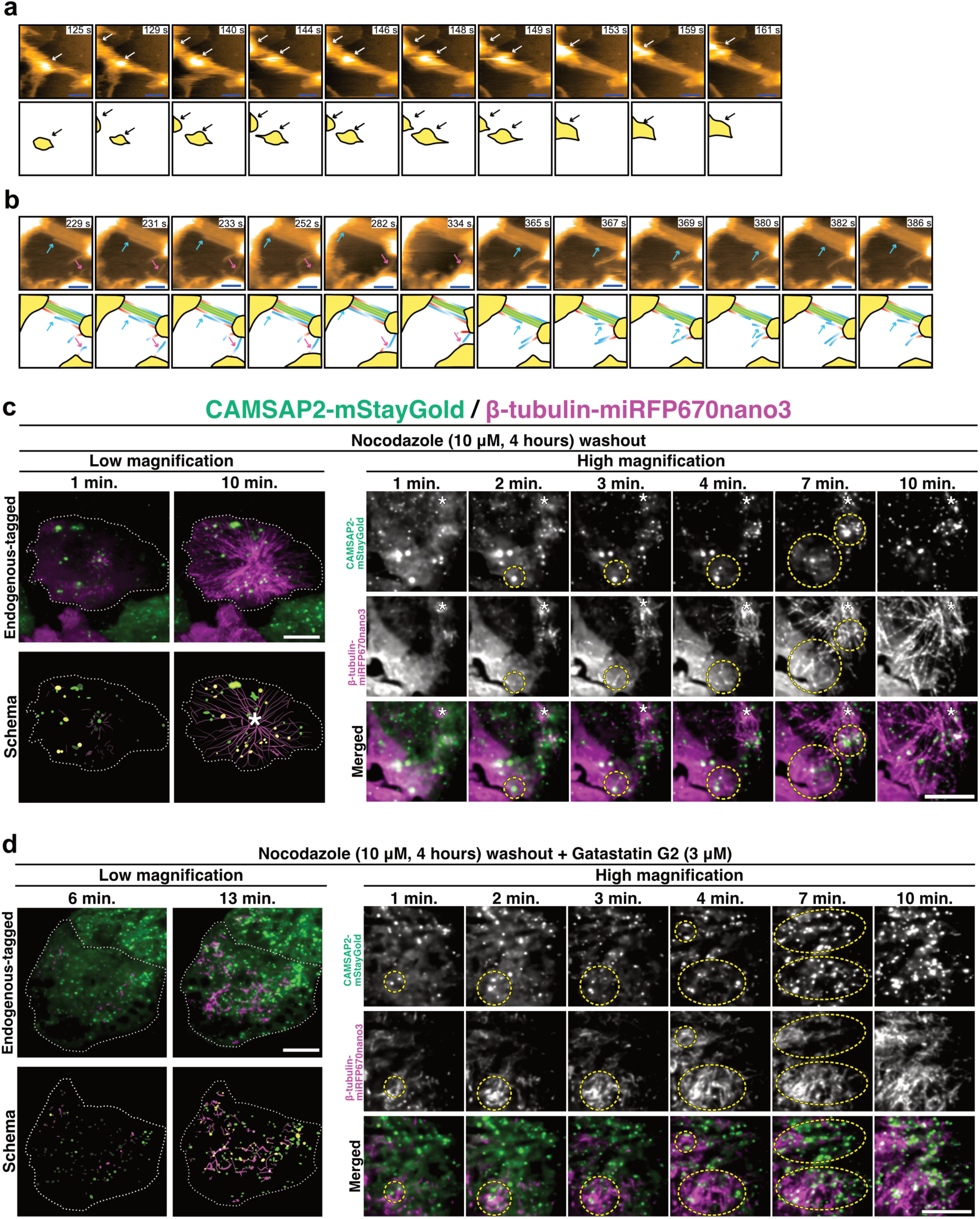
CAMSAP2 defines a γ-TuRC-independent ncMTOC in cells and drives Cam2-aster network formation in cell. **a.** hsAFM images showing Cam2-aster structures. Fusion of two Cam2-asters and the corresponding schematic are shown. **b.** Cam2-aster with branching microtubules extending toward a neighboring aster center (cyan and magenta arrows). Dynamic microtubules with distinguishable polarity are colored red at the plus end and light blue at the minus end, whereas stable microtubule bundles are colored light green. Tubulin and CAMSAP2 CC1–CKK concentrations were 20 µM and 2 µM, respectively. Scale bars, 200 nm. Imaging speed, 1.88 s/frame (see also **Supplementary Video 6**). **c.** Microtubule regrowth in HeLa cells carrying endogenous CAMSAP2–mStayGold (green) and β-tubulin–miRFP670nano3 (magenta) knock-ins. Cells were treated with 10 μM nocodazole for 4 h and imaged from the basal side immediately after washout for 10 min at 1-min intervals. Left, basal images at 1 and 10 min after washout. Asterisks indicate centrosomes. Right, time-lapse images of microtubule regrowth; dashed circles indicate CAMSAP2-dependent microtubule networks. Scale bar, 10 μm. See also **Supplementary** Fig. 11a**,b** and **Supplementary Video 7-9**. **d.** Microtubule regrowth in CAMSAP2–mStayGold/β-tubulin–miRFP670nano3 knock-in HeLa cells after co-treatment with 10 μM nocodazole and 3 μM Gatastatin G2 for 4 h. Imaging was performed immediately after washout for 10 min at 1-min intervals in the continued presence of 3 μM Gatastatin G2. Left, basal images at 1 and 10 min after washout. Right, time-lapse images of microtubule regrowth. Dashed circles indicate regions of interest. Scale bar, 10 μm. See also **Supplementary** Fig. 9c**, 1d, Supplementary Video 10 and 11**.

hsAFM further revealed a stepwise mode of network assembly. Microtubules first emerged from individual condensates and dynamically explored the surrounding space. When a growing microtubule encountered a neighboring condensate, its plus end stabilized, while additional microtubules continued to emerge from both centers (**Fig. 4b**). This process progressively linked multiple condensates into a continuous network.

Because CAMSAP2 binds microtubule minus ends, this geometry implies mixed polarity within connecting bundles, with minus ends anchored at the originating condensate and plus ends extending toward the opposing condensate. As assembly progressed, rings and curved protofilaments diminished, consistent with their incorporation into elongating microtubules (**Supplementary Fig. 10a**). We also observed discrete oligomeric particles moving along bundles between neighboring condensates. One particle traveled ∼500 nm in ∼10 s from one Cam2-aster to another, and some trajectories were bidirectional (**Supplementary Fig. 10b,c**).

Although the identity of these particles remains unresolved, their behavior is consistent with CAMSAP2–tubulin assemblies translocating along microtubules.

Together, these observations demonstrate that CAMSAP2 condensates function as active organizers of microtubule networks in vitro, driving aster formation, fusion, and interconnection through dynamic filament growth.

### CAMSAP2 condensates define γ-TuRC-independent ncMTOCs in cells

To determine whether the CAMSAP2-dependent nucleation mechanism defined in vitro also operate in cells, we generated HeLa cells carrying CRISPR knock-ins of β-tubulin–miRFP670nano3 and CAMSAP2–mStayGold (**Fig. 4c**). The tagged CAMSAP2 recapitulated endogenous localization patterns, enabling direct visualization of microtubule regrowth. (**Supplementary Fig. 9**) ^51–53^.

After nocodazole washout, microtubule regrowth in control cells was dominated by a centrosomal aster, consistent with γ-TuRC-dependent nucleation (**Fig. 4c, Supplementary Fig. 11a,b, Supplementary Video 7-9**). However, time-lapse imaging also revealed peripheral regrowth sites away from the centrosome (**Fig. 4c**, yellow dashed circles). These sites coincided with CAMSAP2-positive condensates and gave rise to local microtubule arrays. Regrowth from these condensates was initiated slightly later than centrosomes, typically 2 to 4 min after washout, whereas centrosomal regrowth began within 1 to 2 min. CAMSAP2 condensates were highly dynamic and often fused, consistent with liquid-like behavior (**Supplementary Fig. 11b, Supplementary Video 9**) ^32^. Microtubule outgrowth from these peripheral condensates preceded CAMSAP2 redistribution toward the centrosomal network, indicating that CAMSAP2 condensates can function as autonomous non-centrosomal microtubule-organizing centers (ncMTOCs) even in the presence of intact γ-TuRC activity.

To test whether these assemblies require γ-TuRC, we inhibited γ-tubulin with Gatastatin G2 ^54,55^. This treatment abolished γ-TuRC-dependent nucleation and centrosomal aster formation, yet microtubule regrowth persisted throughout the cytoplasm, originating from dispersed CAMSAP2 condensates (**Fig. 4d, Supplementary Fig. 11c,d, Supplementary Video 10, 11**). The resulting network formed a fine, cell-wide meshwork distinct from radial centrosomal arrays. These observations show that CAMSAP2 condensates function as γ-TuRC–independent ncMTOCs in cells. Thus, CAMSAP2-based condensates coexist with centrosomal MTOCs under normal conditions and can dominate microtubule organization when γ-TuRC activity is suppressed.

## DISCUSSION

Our findings establish a unified framework in which spontaneous microtubule nucleation, visualized at molecular resolution in vitro, directly underlies non-centrosomal microtubule organization in cells (**Fig. 5**). We identify phase-separated CAMSAP2 condensates as a γ-TuRC–independent microtubule-organizing centers (ncMTOCs) and reveal how a minus-end MAP couples tubulin condensation, oligomer remodeling, and early filament stabilization to drive microtubule network assembly. In this view, CAMSAP2 does not simply act downstream of nucleation but actively reshapes the intrinsic assembly landscape of tubulin to generate productive nuclei. More broadly, these results expand the conceptual framework of microtubule biogenesis beyond a strictly γ-TuRC-centered view and suggest that MAPs can actively convert structurally heterogeneous tubulin intermediates into elongation-competent seeds

**Fig. 5.**
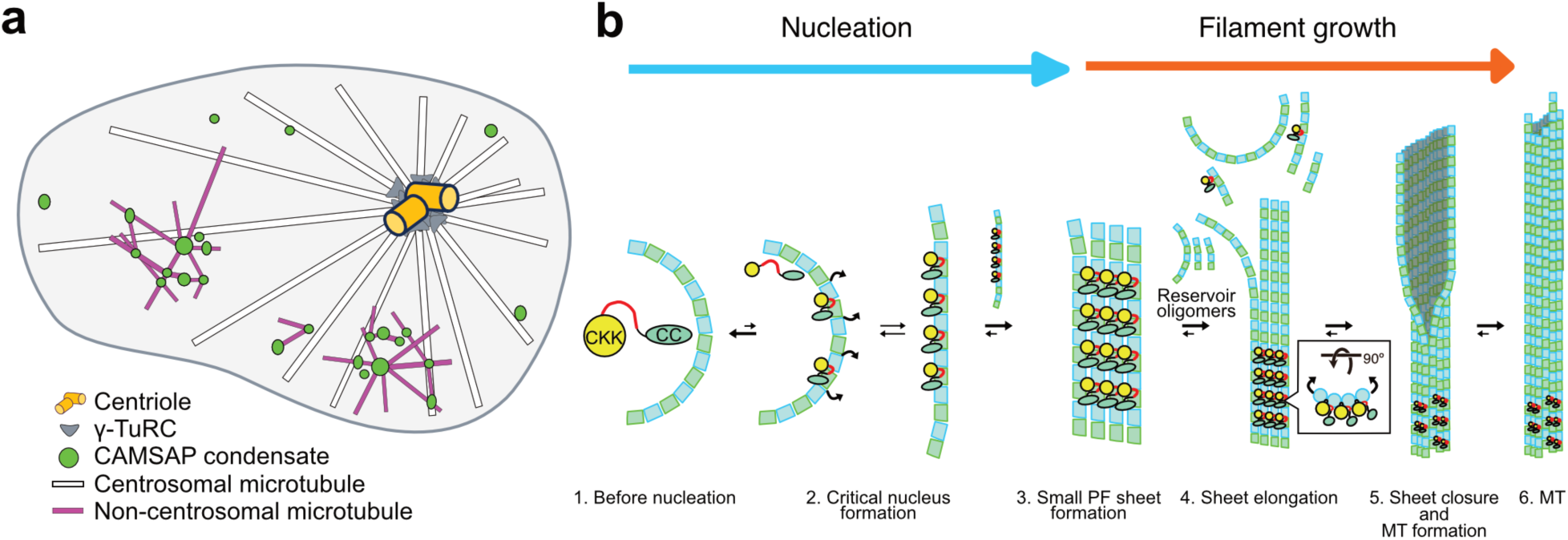
Model of CAMSAP2-driven spontaneous microtubule nucleation at ncMTOCs. **a.** Schematic illustrating a CAMSAP2 condensate-dependent ncMTOC and its associated microtubule network, shown in comparison with the centrosomal MTOC. **b.** Mechanistic model of spontaneous microtubule nucleation driven by CAMSAP2 within condensates (green ovals in panel A).

### CAMSAP2 lowers the energetic barriers of spontaneous nucleation

A central conclusion of this study is that CAMSAP2 directly lowers the two major energetic barriers of spontaneous microtubule nucleation. Classical models propose that nucleation requires the conversion of curved ring-like or short oligomeric tubulin assemblies into straighter protofilaments, followed by the formation of lateral contacts that support sheet formation and tube closure ^7–10,17,19^. Our cryo-EM, hsAFM, MD simulations, and mutational analyses now provide a structural mechanism for this transition. The N-hook engages the tubulin intra-dimer interface and promotes straightening of curved protofilament-like intermediates, whereas the CKK domain stabilizes lateral interactions and favors sheet-like assemblies compatible with 13-protofilament microtubules ^36,37^. CAMSAP2 condensates further potentiate these reactions by concentrating tubulin and enriching oligomeric intermediates through LLPS ^32^. Together, these activities transform a heterogeneous pool of transient assemblies into elongation-competent seeds.

This mechanism revises the long-standing view that ring-like tubulin assemblies are primarily dead-end or depolymerization-associated byproducts ^8,9,18,56,57^. Our observations suggest that closed rings must first open before they can be incorporated into growing microtubules (**Fig. 2**, **Fig. 3d**), consistent with the report by Diaz et al. that closed rings can act as inhibitory species under some conditions ^18^. Rather than bypassing the intrinsic assembly landscape of tubulin, CAMSAP2 exploits and reshapes it to drive nucleation (**Fig. 5b**).

### Oligomer composition encodes early growth dynamics

Our results further indicate that the structural composition of the local oligomer pool governs the earliest stages of microtubule growth. hsAFM imaging on lipid-supported membranes revealed that elongation proceeds discontinuously, with bursts of growth interspersed with pauses and brief shortening events. These dynamic transitions correlate with the structural state of nearby intermediates: straighter protofilament fragments accompany growth, whereas curved oligomers and ring-like species accumulate near paused or shortening ends.

This finding provides a mechanistic link between spontaneous nucleation and early dynamic instability. Rather than emerging only after a mature lattice has formed, pause–growth transitions appear to be encoded already within the heterogeneous oligomer ensemble present at the growing end ^1,13,14,18^. In this view, early elongation is not simply the linear addition of tubulin dimers, but a selective incorporation process in which local oligomer geometry biases end behavior. Because CAMSAP2 selectively accumulates at free minus ends in cells ^28–31^, these mechanisms are likely to be particularly relevant to nucleation-proximal and early elongation events within non-centrosomal arrays.

### CAMSAP2 couples spontaneous nucleation to minus-end organization

The cell-based data show that this molecular mechanism is sufficient to generate bona fide ncMTOC behavior in vivo. Endogenous CAMSAP2 condensates act as autonomous sites of microtubule regrowth, and their activity persists even when γ-TuRC function is suppressed (**Fig. 4**), demonstrating that CAMSAP2-driven assemblies can operate independently of the canonical centrosomal pathway ^1–6,32^. During regrowth, CAMSAP2 first marks dispersed non-centrosomal nucleation sites and associates with newly growing microtubules, then progressively redistributes as local networks merge into the broader array. These dynamics indicate that CAMSAP2 not only initiates local assembly but is also incorporated into the evolving microtubule architecture.

This framework helps reconcile our findings with recent work emphasizing a post-nucleation role for CAMSAP family proteins. Rai et al. showed that CAMSAP2/3 can bind newly formed minus ends of γ-TuRC-nucleated microtubules and promote their release from γ-TuRC, thereby generating free non-centrosomal minus ends ^33^. Rather than conflicting with a nucleation-promoting role, our results suggest that these activities lie along a continuous pathway. CAMSAP2 appears to stabilize structurally fragile early minus-end intermediates, promote their elongation, and subsequently organize them into persistent non-centrosomal arrays. We therefore propose that CAMSAP2 functions not as a simple de novo nucleator, but as a factor that continuously supports nucleation, immediate post-nucleation maturation, and minus-end organization.

### CAMSAP2 defines an integrated, phase-separated nucleation system

Several MAPs, including XMAP215/chTOG, TPX2, CLASP, p150^Glued^, doublecortin, and GAS2-family proteins, have been implicated in lowering the nucleation barrier by stabilizing early intermediates, promoting polymerization, or bridging protofilaments ^4,5,20,24–27^. These activities are often partitioned across multiple factors and, in many cases, have been interpreted as stabilizing or amplifying early nuclei rather than integrating all steps of nucleation and early elongation within a single molecule. CAMSAP2 appears distinct in that it brings several of these activities together within a single LLPS-forming molecule. Within condensates, CAMSAP2 concentrates tubulin, engages oligomeric intermediates, promotes protofilament straightening, stabilizes lateral interactions, and remains associated with the minus end after nucleation^28,32,36,37^.

This mechanism contrasts sharply with γ-TuRC-mediated nucleation, which relies on a preassembled structural template, and may also distinguish CAMSAP2 from many previously described MAP-assisted nucleators ^1–3^. CAMSAP2 does not appear to provide a rigid preassembled template, but instead promotes nucleation by reorganizing endogenous tubulin assemblies and biasing their progression toward productive sheet and tube geometries within a phase-separated environment. Such a mechanism is well suited to dispersed ncMTOCs and may help explain how non-radial microtubule networks are generated in differentiated cells ^4–6,28–31,34,35^.

More broadly, our results suggest that microtubule nucleation can arise from a self-organizing, phase-separated system in which multiple MAP-like activities are spatially concentrated and functionally integrated. This provides an alternative to template-based nucleation and offers a conceptual framework for understanding how diverse microtubule architectures are generated in cells lacking dominant centrosomal control.

### Condensate-based ncMTOCs broaden the design principles of microtubule architecture

An important implication of our findings is that cells may deploy γ-TuRC-dependent and γ-TuRC-independent pathways in parallel to build distinct microtubule architectures. Centrosomal γ-TuRC typically produces radial arrays from a centralized site, whereas CAMSAP2 condensates generate networks from multiple dispersed origins (**Fig. 5a**) ^4–6,28–31,34,35^. These two modes are therefore unlikely to be merely redundant; instead, they encode fundamentally different geometric and mechanical properties of the cytoskeleton.

Multi-origin assembly from dispersed condensates is expected to promote a more homogeneous distribution of filaments, increased architectural flexibility, and enhanced robustness to local perturbations. In contrast to radial arrays, which concentrate forces along defined axes, mesh-like networks assembled from multiple nucleation sites may distribute mechanical stress more evenly and maintain structural integrity under deformation. Because growth is initiated at many sites, local damage may be compensated by continued assembly from neighboring condensates, enabling robust maintenance and repair of microtubule networks. Such properties are likely to be particularly advantageous in polarized or highly differentiated cells, where microtubule organization must be maintained far from the centrosome and dynamically remodeled in response to local cues ^28–31,34,35^.

Importantly, these observations raise the possibility that cell-type–specific microtubule architectures emerge not from a single nucleation mechanism, but from the regulated balance between centrosomal and non-centrosomal pathways. In this context, CAMSAP2 may function not only as a generator of non-centrosomal arrays but also as a modulator of centrosome-derived networks, for example by capturing, stabilizing, or redistributing newly formed microtubules. Such dual functionality would enable cells to integrate radial and mesh-like architectures into a unified cytoskeletal system, providing a versatile strategy for shaping diverse cellular morphologies.

### A cross-scale view of spontaneous microtubule nucleation

Finally, this work establishes an experimental framework for directly interrogating early microtubule assembly intermediates across spatial and temporal scales. By integrating cryo-EM, cryo-ET, hsAFM, MD simulations, and targeted perturbation of the N-hook, we captured ring-to-sheet and sheet-to-tube transitions and linked these intermediates to functional assembly states. This cross-scale approach provides, to our knowledge, one of the clearest direct visualizations of spontaneous microtubule nucleation at near-molecular resolution.

### Concluding remarks

Taken together, our findings revise the prevailing γ-TuRC-centric model of microtubule biogenesis by establishing spontaneous nucleation as a physiologically deployed mechanism underlying non-centrosomal microtubule organization. We show that phase-separated CAMSAP2 condensates function as γ-TuRC–independent microtubule-organizing centers that reshape tubulin oligomers to drive nucleation and early minus-end elongation.

More broadly, these results suggest microtubule network architecture, and thereby cell shape, may be governed by the relative deployment and integration of centrosomal and non-centrosomal nucleation systems. Regulation of tubulin oligomer geometry and composition within condensates, together with their interplay with centrosomal pathways, may represent a general mechanism by which MAPs coordinate nucleation, early growth, and the emergence of diverse cellular morphologies.

## Acknowledgments

We thank K. Chin and N. Shimizu for research management support, M. Takeichi, M. Kikkawa at the University of Tokyo, and the members of the Nitta and Fukuma labs for valuable discussions. The authors express their gratitude to the WPI Nano Life Science Institute (WPI-NanoLSI), Kanazawa University, for facility access and financial support, and to Toshio Ando for instrumental assistance. During the preparation of this work, the authors used ChatGPT to improve writing.

## Funding

Japan Society for the Promotion of Science (JSPS) KAKENHI grants JP21H05254 (RN)

Japan Society for the Promotion of Science (JSPS) KAKENHI grants JP23K24057 (RN)

Japan Society for the Promotion of Science (JSPS) KAKENHI grants JP22K06809 (TI)

Japan Society for the Promotion of Science (JSPS) KAKENHI grants JP25H02379 (TI)

Japan Society for the Promotion of Science (JSPS) KAKENHI grants JP23KJ0472 (HL)

Japan Society for the Promotion of Science (JSPS) KAKENHI grants 21H05251 to (TF)

Japan Society for the Promotion of Science (JSPS) KAKENHI grants 21H05249 to (YS)

Japan Society for the Promotion of Science (JSPS) KAKENHI grants 23K05713 (KXN)

Japan Society for the Promotion of Science (JSPS) KAKENHI grants 23H02452-01 (KXN)

JST Moonshot Research and Development Program grant JPMJMS2024-7 (RN) JST FOREST JPMJFR214K (TI)

Japan Agency for Medical Research and Development (AMED) CREST grant JP21gm161003 (TI)

The Hyogo Science and Technology Association (RN)

Japan Synchrotron Radiation Research Institute (JASRI) (Proposal Nos. 2021B2536, 2022A2536, 2022B2542, 2023A2542, 2023B2537, 2024B2531, and 2024B2532)

AMED-BINDS (JP25ama121001).

## Author contributions

Conceptualization: AY, TI, KXN, TF, RN

Data curation: AY, TI, RK, KXN, TY, HL, HS, MFN, TK

Investigation: AY, TI, RK, KXN, SK, HL, TK

Funding acquisition: TI, RXN, TF, RN

Supervision: TI, YS, TF, RN

Writing – original draft: TI

Writing – review & editing: AY, TI, KXN, TF, RN

## Data and materials availability

Cryo-EM maps of the polymerizing microtubule, microtubule with CAMSAP2-CC3-CKK with 13PF microtubule or 14PF microtubule have been deposited in the Electron Microscopy Data Bank (EMDB) with accession codes EMDB: EMD-67789 and EMDB: EMD- 67790 respectively. Atomic models of the CAMSAP2-CC3-CKK with 13PF microtubule or 14PF microtubule have been deposited in the Protein Data Bank (PDB) with accession codes PDB: 21LB, and PDB: 21LC, respectively. Code for the curvature estimation analyses are openly available in the following GitHub repository: [https://github.com/NewazFahim/Microtubule]. Any data reported in this paper will be shared by the lead contact upon request. Plasmids generated in this study will be shared upon request.

## Supplementary Materials

Materials and Methods

Supplementary Fig. 1 to 11

Tables S1 to S3

References 1-57

Supplementary Video 1 to 11

