## Supplementary Materials for "CAMSAP2 condensates drive a γ-TuRC-independent pathway for non-centrosomal microtubule nucleation"

**Materials and Methods**

### Materials

| REAGENT OR RESOURCE | SOURCE | IDENTIFIER |
|---|---|---|
| **Bacterial and virus strains** | | |
| *E.coli* BL21(DE3) | Novagen | |
| *E.coli* DH5α | | |
| *E.coli.* Stbl3 | Invtirogen | |
| **Biological samples** | | |
| Sus scrofa brain | Kobe process meat co. | N/A |
| **Chemicals, peptides, and recombinant proteins** | | |
| Benzamidine | nacalai tesque | cat # 04036-72 |
| Dithiothreitol (DTT) | nacalai tesque | cat # 14112-94 |
| EDTA | nacalai tesque | cat # 15111-45 |
| HEPES | nacalai tesque | cat # 17514-15 |
| Imidazole | nacalai tesque | cat # 19004-35 |
| Isopropyl-β-D-thiogalactopyranoside (IPTG) | nacalai tesque | cat # 19742-94 |
| KCl | nacalai tesque | cat # 28514-75 |
| Leupeptin | nacalai tesque | cat # 20454-76 |
| $MgCl_2$ | nacalai tesque | cat # 20909-55 |
| NaCl | nacalai tesque | cat # 31319-45 |
| PMSF | nacalai tesque | cat # 06297-31 |
| Pepstatin A | nacalai tesque | cat # 26436-52 |
| Tris-HCl | FUJIFILM | cat # 201-06273 |
| GTP | nacalai tesque | cat # 08921-84 |
| GDP | nacalai tesque | cat # 17354-02 |
| PIPES | DOJINDO | cat # 345-02225 |
| GMPCPP | Jena Bioscience | cat # NU-405L |
| 5(6)-carboxytetramethylrhodamine succinimidyl ester (TMR) | Santa Cruz Biotechnology | cat # sc-495645 |
| N-[6-(biotinamide)hexanoyl]-6-aminohexanoic acid N-succinimidyl ester (Biotin-LC-LC-NHS) | Tokyo Chemical Industry | cat # S0956 |
| DPPC (1,2-dipalmitoyl-sn-glycero-3-phosphocholine) | Avanti Polar Lipids | cat # 850355 |
| DPTAP (1,2-dipalmitoyl-3-trimethylammonium-propane) | Avanti Polar Lipids | cat # 890870 |
| Carbenicillin Disodium Salt | nacalai tesque | Cat#07129-14 |

| | | |
|---|---|---|
| Ampicillin | nacalai tesque | |
| Spectinomycin Dihydrochloride n-Hydrate | nacalai tesque | 32147-81 |
| LB Broth, Miller | nacalai tesque | 20068-04 |
| Taxol | | |
| Nocodazole | | |
| Gatastatin G2 | Funakoshi | Cat # FDV-0040 [1] |
| Dimethyl sulfoxide | FUJIFILM | cat # 045-24511 |
| KOD One PCR Master Mix -Blue- | TOYOBO | Cat # KMM-201X5 |
| NEBuilder HiFi DNA assembly kit | NEB | Cat # E2621 |
| MluI-HF | NEB | cat # R3198S |
| SapI | NEB | cat # R0569S |
| Alkaline Phosphatase (*E.coli* C75) | Takara Bio | Cat # 2120A |
| Ligation High Ver.2 | TOYOBO | Cat # LGK-201 |
| DMEM(4.5g/l Glucose) without L-Gln and Sodium Pyruvate, liquid | nacalai tesque | Cat # 08488-55 |
| Fetal bovine serum | Gibco | Cat # |
| Penicillin-Streptomycin Mixed Solution | Nacalai tesque | Cat # 26253-84 |
| HEPES (1 M) | Gibco | Cat# 15630080 |
| GlutaMAX Supplement | Gibco | Cat# 35050061 |
| Sodium Pyruvate | Nacalai tesque | Cat # 06977-34 |
| TrypLE Express Enzyme (1X), no phenol red | Gibco | Cat # 12604021 |
| Opti-MEM I Reduced Serum Media | Gibco | Cat # 31985070 |
| G418 Disulfate | Nacalai tesque | Cat # 08973-14 |
| Blasticidin S Hydrochloride | FUJIFILM | Cat # 029-18701 |
| HBSS (10X), calcium, magnesium, no phenol red | Gibco | Cat # 14065056 |
| Paraformaldehyde EMPROVE_ ESSENTIAL DAC | Merck | Cat # 104005 |
| Calcium Chloride | NacalaiTesque | Cat # 06729-55 |
| Polyoxyethylene Sorbitan Monolaurate (Tween 20) | NacalaiTesque | Cat # 35624-15 |
| Polyethylene Glycol Mono-p-isooctylphenyl Ether (Triton X-100) | NacalaiTesque | Cat # 12967-45 |
| Skim milk powder | Morinaga Milk Industry Co., Ltd. | N/A |
| FluorSave Reagent | Millipore | Cat # 345789 |
| Anti-α-tubulin mouse monoclonal antibody, clone 12G10 | Developmental Studies Hybridoma Bank | 12G10 anti-alpha-tubulin was deposited to the DSHB by Frankel, J. / Nelsen, E.M. (DSHB Hybridoma Product |

| | | |
|---|---|---|
| | | 12G10 anti-alpha-tubulin) |
| Anti-CAMSAP2 rabbit polyclonal antibody | proteintech | Cat # 17880-1-AP |
| Anti IgG(H+L), Rabbit (Donkey) CF 594 | Biotium | Cat # 20152-1 |
| Anti IgG(H+L), Mouse (Donkey) CF 647 | Biotium | Cat # 20046-1 |
| 4',6-diamidino-2-phenylindole (DAPI) | | |
| **Deposited data** | | |
| PDB: 21LB, 21LC | this paper | |
| EMDB: EMD-67789, EMD-67790 | this paper | |
| **Experimental models: Cell lines** | | |
| Sf9 | Thermo Fisher Scientific | B82501 |
| HeLa cells | JCRB Cell Bank | cat# JCRB9004, lot# 08162018, RRID:CVCL_0030 [2] |
| **Recombinant DNA** | | |
| pFastBac1-His-Strep2-SfGFP-CAMSAP2-FL | Our laboratory [3] | RN189 |
| pET28-His-Strep2-SfGFP-CAMSAP2-CC1-CKK (696-1472) | This study | RN290 |
| pGFPS1-HAT-GFPS1-TEV-CAMSAP2-CC3-CKK (1157-1454)-7xHis | Our laboratory [3] | RN64 |
| pET28-His-Strep2-SfGFP-CAMSAP2-CC3-N-hook (1157-1331) | This study | IMA1432 |
| pFastBac1-His-Strep2-SfGFP-CAMSAP2-3A | This study | RN278 |
| pFastBac1-His-Strep2-SfGFP-CAMSAP2-5A | This study | RN279 |
| pX330S-2-PITCh | pX330S-2-PITCh was a gift from Takashi Yamamoto (Addgene plasmid # 63670 ; http://n2t.net/addgene:63670 ; RRID:Addgene_63670) | RN423 ; Suzuki KT, Yamamoto T. *Nat Protoc.*, 2016. 5/11/26 9:56:00 AM |
| sghtubb4b-C_ pX330S2-PITch | This Study | RN416 [4] sgRNA sequences, CCTAGAGCCTTCAGTCACTG |
| sghCAMSAP2-c_ pX330S-2-PITCh | This study | sgRNA sequences ACCCACTAAGGCATAGAAGT; RN422 |

| | | |
|---|---|---|
| mStayGold-2A-Nw cDNA | This study | Ando et al., Nat. Methos, 2024; E182D [5] |
| pCANw-sal-EGFP (within a Neo(E180D) gene) | Gift from Dr. Masatoshi Takeichi. | RN419 [6] |
| IRES-Neo(weak)_ pBlueScript II-SK(+) | This Study | RN409; E182D [5] |
| qTAG-C-miRFP670nano3-Blast | qTAG-C-miRFP670nano3-Blast was a gift from Laurence Pelletier (Addgene plasmid # 207713 ; http://n2t.net/addgene:207713 ; RRID:Addgene_207713) | RN425 [4,7]; |
| pBlueScriptII-SK(-)-2XPITCh-GOI (IH) | This study | RN424 |
| hCAMSAP2-mStayGold-2A-Nw_pBlueScriptII-SK(-)-2XPITCh | This study | RN420 |
| htubb4B-mStayGold-2A-Nw_ pBlueScriptII-2XPITch | This study | RN418 |
| hTUBB4B-miRFP670nano3-2A-B_pBlueScriptII SK(-)-2XPITCh | This study | RN421 |
| **Software and algorithms** | | |
| ChimeraX v1.9 | Goddard et al. [8] | https://www.cgl.ucsf.edu/chimerax/ |
| Coot v0.9.8.1 | Emsley et al. [9] | https://www2.mrc-lmb.cam.ac.uk/personal/pemsley/coot/ |
| CTFFIND4 | Rohou and Grigorieff, [10] | https://grigoriefflab.umassmed.edu/ctffind4 |
| Microtubule Relion-based pipeline (MiRP) v2 | Cook et al. [11,12] | https://github.com/moores-lab/MiRPv2 |
| SerialEM 3.8 | Mastronarde, [13] | https://bio3d.colorado.edu/SerialEM/ |
| RELION 5.0 | Burt et al. [14] | https://www3.mrc-lmb.cam.ac.uk/relion |
| iMOD 5.1 | Mastronarde and Held, [15] | https://bio3d.colorado.edu/imod/ |
| Phenix v1.21 | Afonine et al. [16] | https://phenix-online.org/ |
| Fiji (Image-J) | Schneider et al. [17] | https://imagej.net/software/fiji/downloads |
| ZEN | Carl Zeiss Microscopy | https://www.zeiss.com/microscopy/en/products/software/zeiss-zen.html |
| NIS-Elements | NIKON Corporation | https://www.microscope.healthcare.nikon.co |

| | | |
|---|---|---|
| | | m/products/software/nis-elements |
| Affinity Designer V2 | Serif Europe Ltd. | https://www.affinity.studio/en |
| Affinity Publisher V2 | Serif Europe Ltd. | https://www.affinity.studio/en |
| UMEX Viewer | In-house software | |
| HS-AFM method | Uchihashi et al., 2012 [18] | https://doi.org/10.1038/nprot.2012.047 |
| **Cryo-EM** | | |
| R1.2/1.3 cupper 300 mesh | Quantifoil | |
| **hsAFM** | | |
| Ultra-short cantilevers for HS-AFM | NANOANDMORE | USC-F1.2-K0.15 |
| **Other** | | |
| Plasmid DNA Extraction Mini Column | FAVORGEN | Cat # FAPDE-C100 |
| FavorPrep Plasmid DNA Extraction Kit | FAVORGEN | Cat # FAPDE 002-1 |
| NucleoBond Xtra Midi | MACHEREY-NAGEL | Cat #740414.50 |
| His-select affinity column | Sigma-Aldrich | H0537 |
| Amicon Ultra concentrator (10 kDa MWCO) | Merck Millipore | UFC8010 |
| Superdex 200 Increase 10/300 GL size exclusion chromatography column | GE Healthcare | 28990944 |
| Superose 6 Increase 10/300 size exclusion chromatography column | GE Healthcare | 29091596 |
| GS4B Sepharose 4 Fast Flow | Cytiva | 17513202 |
| Super Electroporator NEPA21 Type II | Nepa Gene Co., Ltd. | https://nepagene.jp/en/products/electroporation-en/nepa21-electroporator |
| NEPA electroporation cuvette 2 mm gap | Nepa Gene Co., Ltd. | Cat # EC-002S |
| 12- or 13-mm diameter cover glass, No.1 | Matsunami Glass | Cat#C012001, #C013001 |
| Slide glass | Matsunami Glass | Cat#S0318 |
| 35-mm diameter glass bottom dishes | Matsunami Glass | Cat#D11530H |
| Violamo Cell Culture Plate (4 Well) | VIOLAMO | Cat # VTC-P4 |
| Cell Culture Dishes, TC-treated surface, 35 mm | IWAKI | Cat # 3000-035 |
| Costar 6-Well cell culture multiple well plate, flat bottom, with lid | CORNING | Cat # 3516 |
| Confocal laser scanning microscope LSM700 | Carl Zeiss Microscopy | RRID: SCR_017377 |

| | | |
|---|---|---|
| Ti2E microscope equipped with a Ti2-LAPP TIRF (total internal reflection fluorescence microscope) system | NIKON CORPORATION | https://www.microscope.healthcare.nikon.com/products/photostimulation-tirf/ti2-lapp |
| Heating stage | TOKAI HIT | Cat # TPi-TIZH26; https://www.tokaihit.com/products/TPi-TIZH26/ |

#### Methods

##### Cloning and construct design

The generation of full-length and N-terminal truncation constructs of CAMSAP2 has been described previously[3]. The CAMSAP2 CC3 region (residues 1138–1223) was PCR-amplified from RN82 and subcloned immediately downstream of the HRV 3C protease cleavage site in the pGEX-6P vector using the NEBuilder HiFi DNA Assembly kit (New England Biolabs) (construct RN198). For genome editing, we used the Precise Integration into Target Chromosomes (PITCh) method and the qTAG system[4,19]. C-terminal microhomology arms (30–40 bp) for human CAMSAP2 or TUBB4B were PCR-amplified from mStayGold-2A-NeoE182D [5,20] or miRFP670nano3-2A -BSD[7], respectively, and inserted into the pBluescript II SK(–)_2XPITCh_GOI (IH) vector via the two MluI sites (constructs RN420 and RN425). Single-guide RNA (sgRNA) target sequences for human CAMSAP2 were designed using the CRISPR design tool in Benchling (Benchling Inc.). sgRNA target sequences for human TUBB4B were taken from a previous study[4]. Each sgRNA oligonucleotide pair was cloned into the pX330-S2-PITCh vector using the two SapI sites (constructs RN416 and RN422). The coding regions of all constructs were verified by Sanger sequencing. All constructs used in this study are listed in the Key Resources Table.

##### Cell Culture

HeLa cells were purchased from JCRB (cat# JCRB9004, lot# 08162018, RRID:CVCL_0030). Cells were grown in a Dulbecco's Modified Eagle Medium high glucose (Nacalai Tesque) supplemented with 10% fetal bovine serum (FBS) (Gibco), 2 mM GlutaMAX supplement (Gibco), 1 mM sodium pyruvate (Nacalai Tesque), 100 μg/ml Penicillin-Streptomycin mixed solution (Nacalai Tesque) (DMEM) at 37ºC with 5% CO2. The identity of the HeLa cell line provided from JCRB has been authenticated by STR profiling and confirmed negative for either bacterial, fungal, or mycoplasma contamination.

##### Generation of endogenous knock-in cells

We collectively used two endogenous knock-in techniques of qTAG [4] with PITCh [19] systems. HeLa cells were transfected with various combinations of expression vectors in 35-mm diameter dishes. Transfection were performed with Super Electroporator NEPA21 Type II (Nepa Gene Co., Ltd.) according to the manufacturer's instructions. Briefly, cells were placed into a NEPA electroporation cuvette 2 mm gap (NEPA GENE Co. Ltd., #EC-002S) that were filled with 200 μL of Opti-MEM (Gibco) containing 5 μg of each donor vector and 5 μg of spCas9 and sgRNA co-expression vectors. The poring pulse was set to voltage: 150 volts, pulse width: 7.5 milliseconds, pulse interval: 50 milliseconds, number of pulses: +2, and decay rate 10%. The first and second transfer pulse were set to voltage: 20 volts, pulse width: 50 milliseconds, pulse interval: 50 milliseconds, number of pulses: ±5, and decay rate 40%. After electroporation, cells were transfer to warm medium at 37ºC with 5% CO2, immediately. To edit the genome DNA sufficiently, transfected cells were cultured for 3 days. Cells were then selected using 200 μg/ml G418 disulfate (NacalaiTesque) or 10 μg/ml blasticidin S hydrochloride (FUJIFILM Wako Pure Chemical Corporation), respectively. Double knock-in cells were confirmed proper localizations by confocal laser scanning microscopes. These multiple clones were maintained in the presence of G418 disulfate and blasticidin S hydrochloride.

##### Immunofluorescence (IF)

Cells cultured for 2–3 days were fixed with 4% paraformaldehyde (PFA; Merck) in Hank's balanced salt solution (HBSS) containing 1 mM $CaCl_2$ and $MgCl_2$ (Gibco) and supplemented with 10 mM HEPES (Gibco) (HBSS CM+) at 37°C for 15 min. After fixation, samples were washed three times with Tris-buffered saline (TBS; pH 7.5) containing 1 mM $CaCl_2$ (Nacalai Tesque) and 0.005% Tween-20 (Nacalai Tesque) (TBS-C), and permeabilized with 0.25% Triton X-100 (Nacalai Tesque) in TBS-C at room temperature for 5 min. Samples were then blocked with 5% skim milk (Morinaga Milk Industry) in TBS-C at room temperature for 30 min, followed by incubation with primary antibodies diluted in 2.5% skim milk in TBS-C at 4°C overnight. After three washes with TBS-C, samples were incubated with secondary antibodies and DAPI diluted in 2.5% skim milk in TBS-C at room temperature for 1 h. Samples were washed three times with TBS-C and mounted on glass slides using FluoroSave reagent (Millipore).

Images were acquired using confocal laser scanning microscopes (LSM700, Carl Zeiss Microscopy) equipped with a Plan-Apochromat 63×/1.4 objective (Carl Zeiss Microscopy). Images were processed and analyzed using ImageJ and ZEN 3.13 software (Carl Zeiss Microscopy).

**Disassembly of the microtubule polymerization on HeLa cells**

Cells were incubated in DMEM supplemented with 10 μM nocodazole for 4 hours at 37ºC with 5% CO2. The cells were then washed 15 ml of warm phosphate-buffered saline (-). Live cell imaging was acquired a minute for a duration of over 15 minutes after nocodazole removal. Optionally, DMEM were supplemented with 3 μM Gatastatin G2.

**Live cell imaging and in vitro microtubule imaging**

Live imaging of mStayGold and miRFP-670nano3 double knock-in HeLa cells was performed using TIRF microscopy at 37°C. The TIRF system consisted of a Ti2E microscope (Nikon Corporation) equipped with a Ti2-LAPP TIRF system (Nikon Corporation), an iXon Life 897 EMCCD camera (Andor Instruments), a CFI Apochromat TIRF 100XC/1.49 lens (Nikon Corporation), and a heating stage (TOKAI HIT). These cells were seeded onto a 35-mm diameter glass bottom dish (Matsunami Glass). The cells were then cultured for another 2-3 days. Before experiments, the cell media were replaced with 2 ml of HBSSCM+ supplemented with 2% FBS, 2 mM GlutaMAX supplement, 1 mM sodium pyruvate, 100 μg/ml Penicillin-Streptomycin mixed solution. These images were processed and analysed with NIS-Elements software into "Denoise.ai" function (Nikon Corporation) and ZEN software into ''bioformat import'' function (Carl Zeiss Microscopy). Optionally, HBSSCM+ were supplemented with 3 μM Gatastatin G2.

**Tubulin preparation for in vitro studies**

Tubulin was purified from porcine brain tissue as described previously [3,21]. Fluorescently labeled tubulin was prepared by labeling purified tubulin with 5(6)-carboxytetramethylrhodamine succinimidyl ester (TMR) (Santa Cruz Biotechnology, Dallas, TX, USA) or N-[6-(biotinamide)hexanoyl]-6-aminohexanoic acid N-succinimidyl ester (Biotin-LC-LC-NHS) (Tokyo Chemical Industry, Tokyo, Japan), as described previously [3].

**Protein expression and purification**

GFP-tagged full-length CAMSAP2 was expressed in a baculovirus expression vector system (BEVS) using insect cells, following our previous protocol [22]. CAMSAP2 deletion constructs were expressed in Escherichia coli strain BL21(DE3) (Novagen). Cells were grown in LB medium supplemented with 25 μg/mL kanamycin at 37 °C to an OD600 of 0.6–0.8, cooled to

18 °C, induced with 0.1 mM IPTG, and cultured overnight. Cells were harvested by centrifugation, and pellets were stored at −80 °C.

Purification of full-length CAMSAP2 constructs expressed in insect cells was performed as described previously [3]. Briefly, cells were lysed in lysis buffer A (50 mM HEPES pH 7.5, 400 mM KCl, 5 mM imidazole, 10% glycerol, 5 mM β-mercaptoethanol) supplemented with a protease inhibitor mix (0.7 µM leupeptin, 2 µM pepstatin A, 1 mM PMSF, and 2 mM benzamidine), and the lysate was clarified by centrifugation. The supernatant was applied to a His-Select affinity column (Sigma-Aldrich) equilibrated in lysis buffer A. After several washes, bound protein was eluted with 300 mM imidazole. The eluate was concentrated using an Amicon Ultra concentrator (Merck Millipore; 10 kDa MWCO) and loaded onto a Superose 6 size-exclusion chromatography column (GE Healthcare). Peak fractions were collected, concentrated, flash-frozen in liquid nitrogen, and stored at −80 °C.
His-tagged constructs expressed in E. coli were stored at −80 °C until purification. Frozen cell pellets were thawed, resuspended in lysis buffer (40 mM Na-phosphate buffer pH 7.4, 400 mM NaCl, 5 mM imidazole, 14 mM β-mercaptoethanol, 0.7 µM leupeptin, 2 µM pepstatin A, 1 mM PMSF, and 2 mM benzamidine), and lysed by sonication. The lysate was clarified by centrifugation (16,000 × g), and the supernatant was applied to a His-Select affinity column equilibrated in lysis buffer A with protease inhibitors. The column was washed with approximately 10 column volumes of lysis buffer and eluted with approximately 4 column volumes of elution buffer containing 300 mM imidazole. The eluate was concentrated using an Amicon Ultra concentrator (10 kDa MWCO) and applied to a Superdex 200 Increase 10/300 GL size-exclusion column (GE Healthcare) equilibrated in sizing buffer (20 mM HEPES -KOH pH 7.5, 300 mM NaCl, 1 mM DTT). Peak fractions were pooled, concentrated with an Amicon Ultra 10 kDa MWCO concentrator, flash-frozen in liquid nitrogen, and stored at −80 °C.

**Spontaneous nucleation assay (pelleting assay)**
Spontaneous nucleation assays were performed as described previously [3]. Briefly, different concentrations of tubulin were prepared in PEM buffer supplemented with 1 mM GTP (PEM-GTP) to test the intrinsic nucleation ability of tubulin. To test the nucleation activity of CAMSAP2, 1 µM CAMSAP2 mutants (CKK, CC3, CC3–CKK) were mixed with varying concentrations of tubulin. The indicated concentrations of tubulin, with or without 1 µM CAMSAP2 mutants, were incubated on ice for 30 min and then at 37 °C for 30 min. At this point, a small aliquot was taken for total-sample analysis by SDS-PAGE. Reaction tubes were then centrifuged in a TOMY MX-307 for 20 min at 15,000 rpm and 35 °C. The supernatant was discarded, and the pellet was washed with warm PEM-GTP buffer and centrifuged again. Pellets were resuspended in cold CAMSAP2 buffer (20 mM HEPES-KOH pH 7.5, 300 mM NaCl, 1 mM DTT) to depolymerize microtubules, incubated on ice for 30 min, and centrifuged for 20 min at 15,000 rpm and 4 °C to remove aggregates or debris. Supernatants were analyzed by SDS-PAGE. Bands from total and depolymerized samples were quantified using a standard curve generated from tubulin aliquots at 1, 2, 3, and 4 µM, using FIJI (ImageJ) software [17].

**Turbidity assay**
Microtubule nucleation and polymerization were measured by optical density at 350 nm [23] using a spectrophotometer (DS-11+, DeNovix). Tubulin and CAMSAP2 solutions were thawed on ice and centrifuged at 20,000 × g for 2 min at 4 °C to remove aggregates. Mixtures containing 8.5 µM tubulin and 0.5 µM of CAMSAP2 CC1-CKK or mutants at the indicated concentrations were prepared in PEM-GTP buffer. Tubulin and CAMSAP2 were mixed in the final step to

minimize complex formation due to transient high local concentrations. The mixtures were immediately loaded into a 10 mm cuvette in a spectrophotometer pre-warmed to 37 °C, and absorbance at 350 nm was recorded for 30 min at 30 sec intervals.

**Microtubule stabilization assay**

Tubulin (50 µM) was polymerized in PEM-GTP buffer at 37 °C for 30 min. A 16 µL solution containing 1.25 µM CAMSAP2 or mutants in PEM-GTP buffer was prepared on ice. Subsequently, 4 µL of polymerized microtubules was added to the CAMSAP2 solution to achieve final concentrations of 10 µM tubulin and 1 µM CAMSAP2 with a final reaction volume of 20 µl at 37 °C. The mixture was incubated on ice for 5 min and then centrifuged in a TOMY MX-307 at 16,000 × g for 20 min at 4 °C. The supernatant was discarded, and the pellet was retained for further analysis. To the pellet, 20 µl cold depolymerization buffer (50 mM HEPES-KOH pH 7.4, 400 mM KCl, 1 mM $MgCl_2$) was added to the pellet, which was resuspended by pipetting, thoroughly mixed, and incubated on ice for 20 min. Samples were centrifuged again in a TOMY MX-307 at 16,000 × g for 20 min at 4 °C. The resulting supernatants were analyzed by SDS-PAGE, and band intensities were quantified using FIJI (ImageJ) [17].

**Negative-stain electron microscopy**

Samples were prepared in PEM buffer (100 mM PIPES pH 6.8, 1 mM $MgCl_2$, 1 mM EGTA, 1 mM GTP) supplemented with 50 mM KCl, adjusted to 10 µM tubulin by adding 1 µl of CAMSAP2 CC1–CKK or its mutants. The mixtures were incubated on ice for 30 min and then at 37 °C for 10 min. After incubation, 4 µl of each sample was applied to a glow-discharged, carbon-coated 200-mesh copper grid (EM Japan). Grids were stained with 2% uranyl acetate, excess stain was blotted with filter paper, and grids were air-dried. Samples were imaged on a JEM-1400 Plus transmission electron microscope (JEOL) operated at an accelerating voltage of 120 kV, and images were recorded with a JEOL Matataki Flash camera at nominal magnifications of 600- and 50,000-fold.

**Cryo-electron tomography**

A holey carbon grid (R1.2/1.3, Cu, 300 mesh; Quantifoil) was glow-discharged (20 Pa, 10 mA, 30 s) and rendered hydrophilic by applying 0.01% NP-40 aqueous solution. A mixture of 30 µM tubulin, 3 µM CAMSAP2 CC1–CKK, 0.01% NP-40, 1 mM GTP, and 10-nm gold fiducials (CGM5K-10-50, Cytodiagnostics) was prepared on ice, incubated at 37°C for 30 s, and 4 µL of the mixture was applied to the grid in a Vitrobot Mark IV (Thermo Fisher Scientific) maintained at 37°C and 100% humidity. Grids were blotted with a blot force of 10 for 3 s and plunge-frozen into liquid ethane. Data were collected on a Glacios transmission electron microscope (Thermo Fisher Scientific) operated at 200 kV and equipped with a Falcon 4i direct electron detector (Thermo Fisher Scientific), using Tomography 5 software (Thermo Fisher Scientific). Tilt series were acquired from −51° to 51° in 3° increments, with a total accumulated dose of ~110 $e^-/Å^2$ and a target defocus range of −2 to −4 µm. A nominal magnification of 45,000× was used, corresponding to a pixel size of 3.1 Å.

Collected movies were processed using the RELION tomography toolkit [14]. Movies were first motion-corrected with odd/even frame splitting and CTF-corrected. After manual exclusion of poor-quality tilt images, tilt series were automatically aligned using gold fiducials. Half tomograms were reconstructed at 7 Å/pixel, and three tomograms containing representative protein densities were used to train a cryo-CARE denoiser [24]. The trained denoiser was then applied to all tomograms, and the correctness of tilt-series handedness was verified at this stage by inspection of the microtubule lattice pattern.

**Grid preparation and cryo-EM data collection for high resolution reconstruction**

Tubulin (40 µM) was polymerized in PEM-GTP buffer at 37 °C for 30 min. A 4 µL aliquot of polymerized microtubules was applied to a glow-discharged carbon grid (R2/2; 200 mesh, Quantifoil). After incubation for 30 s at 37 °C and 90% humidity in a Vitrobot Mark IV (Thermo Fisher Scientific), excess solution was blotted away with Whatman no. 1 filter paper. Subsequently, 4 µL of 40 µM CAMSAP2 CC3–CKK was added to the grid. After a further 30 s incubation, the grid was blotted for 5 s with a blot force of 5 and plunge-frozen into liquid ethane. Data collection was performed on a 200 kV Glacios TEM (Thermo Fisher Scientific) equipped with a K2 Summit detector (Gatan) at 36,000× magnification under low-dose conditions, controlled by SerialEM software [13] at SPring-8. All data were collected as 40-frame movie stacks, with a total electron dose of 50 $e^{-}/Å^2$ and a pixel size of 1.15 Å/pixel. The defocus range was set between −2.5 and −1.0 µm.

**Image processing and 3D reconstruction from cryo-EM images**

For high-resolution cryo-EM single-particle analysis of the CC3–CKK–microtubule complex, we used a RELION-based image-processing pipeline for pseudo-helical microtubules (MiRP ver. 1) [11,12]. Microtubules were manually picked and segmented with a separation distance of 82 Å (approximately the tubulin dimer repeat) using the helical mode in RELION ver. 3.0.8 [25] Particles were extracted with 4× binning and averaged over seven adjacent segments to generate high signal-to-noise super-particles. Low-pass-filtered 3D models of microtubules with 11–16 protofilaments were used as references for initial classification. The dominant 13- and 14-protofilament microtubules were selected for further analysis.

Initial rotational alignment was performed with a wide search, followed by refinement with tightly restrained Psi and Tilt angles and broader searches for rotational and translational shifts. To improve alignment accuracy, the most frequently assigned rotational angle for each microtubule was determined and applied. Misassignments of X/Y shifts were detected by plotting shift values against particle indices and corrected by adding or subtracting integer twists to remove abrupt jumps caused by α/β-tubulin similarity. After refining X/Y shifts, supervised 3D classification was carried out using references for all possible seam positions, including counterparts shifted by 41 Å along the helical axis. Unified classes were re-extracted without binning and refined with or without helical symmetry for high-resolution reconstruction. Bayesian polishing and CTF refinement were then performed, and refined maps were post-processed using automatic B-factor estimation in RELION. Final resolutions were determined using the gold-standard Fourier shell correlation at the 0.143 criterion (Table S1).

The structural model was built by fitting PDB ID 6QUS [26] into the density using Chimera [27], ChimeraX[28] replacing the sequence with that of CAMSAP2 in COOT [9], and refining using Phenix [29]. The N-hook region was manually modeled in COOT.

**Preparation of lipid bilayer**

Liposomes composed of 1,2-dipalmitoyl-sn-glycero-3-phosphocholine (DPPC; Avanti Polar Lipids, Alabaster, AL) and 1,2-dipalmitoyl-3-trimethylammonium-propane (DPTAP; Avanti Polar Lipids) first optimized for lipid ratio at a 92.5/7.5 (wt/wt) ratio, and mica-supported lipid bilayers, were prepared according to our previous protocol [30]. This positively charged lipid bilayer was used to gently immobilize tubulin and to observe rings, protofilaments, andmicrotubule intermediate structures, including tubulin sheets, mature microtubules, and aster structures, in the presence or absence of CAMSAP2 on membrane surface during hsAFM experiments. An additional advantage of using this lipid membrane bilayer was that non-specific bindings of CAMSAP2 were not observed, ensuring nearly all proteins were involved in the reaction in solution or at the membrane interface.

**hsAFM Measurements**

AFM experiments were performed using a custom-built high-speed atomic force microscope (hsAFM) developed by Ando and colleagues [31]. hsAFM operated in intermittent tapping mode with amplitude modulation feedback in solution, using ultra-short cantilevers (USC-F1.2-K0.15; NANOANDMORE) with a spring constant of approximately 0.15 N/m, a resonant frequency of approximately 1.2 MHz, and a quality factor of approximately 3.0 in air. hsAFM imaging was carried out essentially as described previously [30,32,33] and the detailed protocols [18], except that we used a newly developed only tracing imaging (OTI) mode [34]. For hsAFM imaging, the peak-to-peak free oscillation amplitude ($A_0$) of the cantilever was set to approximately 1.6–1.8 nm, and the feedback setpoint was adjusted to approximately 0.9 $A_0$. AFM image rendering and data processing were performed using custom-made analysis software, including Kodec and UMEX Viewer, as described previously [35].

**Preparation of tubulin and CAMSAP2 proteins for hsAFM**

Stock tubes of tubulin and CAMSAP2 (wild-type or mutant constructs CC1–CKK, N-hook, CC1–CKK 3A, and CC1–CKK 5A) stored at –80 °C were thawed on ice and clarified by micro–ultracentrifugation (Himac CS100GXII, Hitachi) at 743,000 × g for 5 min at 4 °C. Clarified supernatants were immediately diluted to the desired working concentrations in cold PEM buffer on ice. Protein concentrations were determined using the Bradford assay (Bio-Rad).

**hsAFM imaging of tubulin rings and protofilaments on mica**

In the imaging chamber, 63 μL of warm PEM buffer (1 mM GTP, equilibrated to 25 °C) was first introduced. Next, 7 μL of 10 μM tubulin in cold PEM buffer was added to reach a final volume of 70 μL containing 1 μM tubulin. A freshly cleaved mica surface mounted on a 2-mm glass rod was firmly glued onto the z-stage of a custom-built hsAFM scanner. The scanner was operated in an inverted configuration, allowing the mica surface to be immersed in the tubulin solution. Using in-house UMEX Sample-scan hsAFM software, the sample surface was automatically approached to the cantilever tip.

Tubulin rings and protofilaments formed in the warm PEM (+GTP) buffer and adsorbed onto the mica surface during this approach, typically within 2–3 min. After confirming their presence (~1 min), 7 μL of 1.1 μM CAMSAP2 or mutant protein in cold PEM buffer was rapidly injected and gently mixed by pipetting 10 times, resulting in a final protein concentration of 0.1 μM. Continuous hsAFM imaging was performed immediately to monitor CAMSAP2-mediated interactions and conformational transitions of tubulin rings and protofilaments on mica.

After preliminary optimization of the acquisition conditions, all hsAFM imaging was conducted at room temperature (~25 °C) using the in-house small hsAFM scanner. Specific scanning sizes, spatial resolutions, and temporal resolutions of AFM images are provided in the corresponding figures and movies.

**hsAFM imaging of tubulin rings, protofilaments, sheets, and microtubules on lipid membranes**

To visualize structures on lipid bilayers composed of DPPC/DPTAP (92.5/7.5, wt/wt), a similar procedure was used, substituting mica with a mica-supported lipid bilayer, prepared as described previously.

In the imaging chamber, 65 µL of warm PEM buffer (1 mM GTP, 25 °C) was added. A freshly cleaved mica-supported membrane mounted on a 2-mm glass rod was fixed to the z-stage of either a small or large custom-built hsAFM scanner operating in inverted configuration. After confirming membrane quality, 6.5 µL of 220 µM tubulin in cold PEM buffer was added and gently mixed by pipetting 10 times, yielding a final 71.5 µL of tubulin concentration of 20 µM.

For control experiments without CAMSAP2, continuous hsAFM imaging was initiated immediately to follow the formation and conformational transitions of protofilaments, rings, sheets, and intermediate or mature microtubules on the membrane surface.

To assess the effects of CAMSAP2 and its mutants on structural transitions (e.g., ring opening or protofilament straightening, sheet formation, microtubule maturation), 2 µM CAMSAP2 or mutant protein was added after 30 min to 1 h of control imaging. This was achieved by rapidly injecting 7.15 µL of 22 µM CAMSAP2 or mutant protein and mixing gently 10 times. Continuous hsAFM imaging then resumed immediately.

All measurements were performed at ~25 °C using the in-house small or large hsAFM scanner. Specific scanning sizes, spatial resolutions, and temporal resolutions of AFM images are provided in the corresponding figures and movies.

**hsAFM scanner specifications**

- **Small hsAFM scanner:**
  Maximum scanning range:
  2,060 × 4,335 × 160 nm (x, y, z)
  Piezoelectric coefficients: 15.92 nm/V (x), 32.5 nm/V (y), 4.0 nm/V (z)
- **Large hsAFM scanner:**
  Maximum scanning range: 8,999 × 6,968 × 770 nm (x, y, z)
  Piezoelectric coefficients: 58.8 nm/V (x), 54.3 nm/V (y), 19.25 nm/V (z)

**Procedure for estimating curvature from hsAFM images**

AFM images were imported into Adobe Illustrator, where structural contours were manually traced to generate vector paths (Supplementary Fig. 5e,f). The coordinates of anchor points along each path were extracted into a text file using custom JavaScript scripts developed in-house and available on GitHub. The extracted coordinate data were then fitted to circular arcs using a least-squares optimization algorithm. The resulting fitted circles are presented in Supplementary Fig. 5e,f. The curvature for each path was calculated as the inverse of the radius of the corresponding fitted circle.

**Procedure for estimating total length of the tubulin fragments from AFM images**

Original AFM images were imported into Adobe Illustrator. Then, we manually drew lines over all segments recognized in the image. Their vector path data was extracted into a text file by custom-made JavaScript. For each segment, the cumulative Euclidean distance between consecutive points was calculated to estimate the arc length. To minimize discretization errors due to uneven point spacing, each path was reparametrized using linear interpolation and uniformly resampled into 1000 equally spaced points along its arc length. Each segment length was then recalculated from the interpolated coordinates as shown in Supplementary Fig. 5f. Finally, the total segments length per image was obtained by summing the lengths of all individual segments.

**Microtubule Curvature Analysis**

Precise curvature calculations for objects in HS-AFM images (or Movies) were achieved through a semi-automated process. Raw images were first imported into Adobe Illustrator, where paths were manually traced over the structures. The number of anchor points on each path was then increased, and their coordinates were extracted into a text file. These tasks were accomplished using two JavaScript scripts developed for Adobe Illustrator's scripting API to automate path manipulation. The resulting coordinate text file was subsequently processed by a Python script that applied a least-squares optimization algorithm to fit circles to the points, from which radii—and thus curvatures—were computed for each path.

**Enhancing the Anchor Points**

The first script utilizes Adobe Illustrator's scripting API to automate the precise subdivision of path segments within vector graphics, streamlining the process of detailed design modifications. Initially, the script checks the version of Adobe Illustrator to ensure compatibility and adjusts its behavior accordingly. It then identifies and extracts path items from the user's selection. Based on a user-defined parameter, the script calculates the positions for new anchor points along the selected segments, which are then inserted to divide the segments into smaller, evenly spaced sections. This automation significantly reduces the manual effort typically required for such tasks, enhancing both the accuracy and efficiency of segment manipulation in vector graphic design workflows. The script's ability to consistently and accurately divide path segments makes it a valuable tool for designers working on complex illustrations, where precision is paramount.

**Extracting the Coordinates of Anchor Points**

The second JavaScript script for Adobe Illustrator automatically collects and saves the positions of path segments from vector graphics. It uses an immediately invoked function expression (IIFE) to keep its scope separate from other scripts. The main function, CreateSegPosition, iterates over all path items in the active Illustrator document and gathers the coordinates of anchor points (the key points that define each path), storing this information in an array. After collecting the data, the script saves it to a text file specified by the user. By using prototype-based methods to efficiently read, store, and write anchor point positions, the script streamlines the extraction of path data, reducing manual work and minimizing errors. Its structure and use of local variables make it reliable and easy to use for documenting and analyzing vector graphic designs in Illustrator.

**Calculating the Curvatures**

This Python script facilitates the analysis of geometric properties of paths extracted from an Adobe Illustrator document by processing anchor point data stored in a text file. The script begins by loading the data from the text file and parsing it into a structured format using pandas and defaultdict. Each path's anchor points are extracted, and the script employs a least-squares optimization algorithm to fit circles to these points. The optimization minimizes the algebraic distance between the data points and the mean circle centered at $(x_c,y_c)$, defined by the equation:

$F_2(c)=R_i-|R|$

where $R_i=\sqrt{(|x-x_c|)^2+(|y-y_c|)^2}$

represents the distance from each point $(x,y)$ to the circle center, and $|R|$ is the mean distance of all points to the center. The radii of the fitted circles are calculated, enabling the determination of curvature for each path. The results are visualized through histograms of the radii and curvatures, as well as a graphical representation of the paths overlaid with their corresponding fitted circles.

This approach provides a quantitative and visual assessment of the curvature characteristics of the paths, aiding in the geometric analysis of vector graphics in Illustrator.

**Computational setup for modeling and simulations of the tubulin complex**

Two simulation systems were constructed: one with CAMSAP2 bound to the central β/α-tubulin dimer and one without CAMSAP2. Atomic coordinates of tubulin and CAMSAP2 were taken from the cryo-EM structure determined in this study. GTP in α-tubulin and GDPin β-tubulin were retained in the nucleotide-binding sites of tubulin subunits, as observed in the density maps. In the β/α dimer, the bottom was supported by β-tubulin and the top by α-tubulin. Residues α193D, α203E, β168D, β205E, and β429D were assigned as protonated based on pKa predictions from PROPKA [36,37]. Missing loop regions in the cryo-EM model were manually built in COOT [9]. Each structure was solvated in a rectangular box with 120 Å×100 Å×210 Å, containing CHARMM TIP3P water molecules [38] and neutralized with 150 mM KCl. Detailed system compositions are summarized in Table S3. The CHARMM c36m force-field parameters [39] were used for tubulins, Guanine nucleotides, and ions. Water molecules are considered as rigid using SETTLE constraints [40]. All bonds involving hydrogen atoms are constrained with SHAKE/RATTLE [41,42].

All MD simulations were performed using the GENESIS software version 2.1 [43]. The simulations were conducted in the NPT ensemble at 300 K and 1 atm, using the stochastic velocity rescaling thermostat and MTK style barostat [44,45] using the refined temperature definition [46]. Group temperature/pressure is used for evaluation of temperature/pressure in thermostat/barostat [47,48]. The nonbonded interactions were calculated using the CHARMM standard procedures: the LJ interaction was smoothly truncated from 10 Å to 12 Å using the force-based switch function [49], and the electrostatic interactions were computed with the particle mesh Ewald summation algorithm [50,51]. For all executions, except for equilibration, we utilized the r-RESPA (or multiple time step) integration with 3.5 and 7.0 fs time steps for the real- and reciprocal-space interactions, respectively [52]. For each system, three independent MD simulations of 1 µs duration were performed in total 3 µs per condition. In one trajectory of the 4tub–Cam2 system, spontaneous dissociation of the N-hook from the tubulin groove was observed and this trajectory was treated as a separate condition (N-hook-dissociated). To compensate for this separation and to maintain comparable sampling for the N-hook–bound condition, an additional independent simulation of the 4tub–Cam2 system was performed, resulting in four trajectories in total for this system.

**Computational analysis**

The structural parameters of tubulin tetramers (protofilament swing, dimer twist, and curvature) were calculated as follows. From each MD trajectory, 50 snapshots were extracted at 20-ns intervals. As illustrated in Supplementary Fig. 6d and 8, the vectors $\boldsymbol{x}_0$, $\boldsymbol{y}_0$ and $\boldsymbol{z}_0$ were defined for tubulin 0, and $\boldsymbol{x}_1$, $\boldsymbol{y}_1$ and $\boldsymbol{z}_1$ for the adjacent tubulin (+1) (Supplementary Fig. 8b), using the centers of mass of the indicated residues (α-tubulin S170 and M415, and β-tubulin S168), computed with the Biopython package [53]. Structural parameters in radians were defined as: protofilament swing = $\arctan(\hat{\boldsymbol{z}}_1 \cdot \hat{\boldsymbol{x}}_0 / \hat{\boldsymbol{z}}_1 \cdot \hat{\boldsymbol{y}}_0)$, dimer twist = $\arctan(\hat{\boldsymbol{y}}_1 \cdot \hat{\boldsymbol{x}}_0 / \hat{\boldsymbol{y}}_1 \cdot \hat{\boldsymbol{y}}_0)$, curvature = $\arcsin(\hat{\boldsymbol{z}}_0 \times \hat{\boldsymbol{z}}_1 / |\hat{\boldsymbol{z}}_0 \times \hat{\boldsymbol{z}}_1|)$, where the hat indicates normalization: $\hat{\boldsymbol{v}} = \boldsymbol{v}/|\boldsymbol{v}|$.

**Quantification and Statistical analysis**

Cryo-EM resolution estimates were performed in Relion using the 0.143 FSC criterion proposed [54]. Validation metrics for atomic models (Table S1) were calculated with Phenix.

#### Supplementary Figures

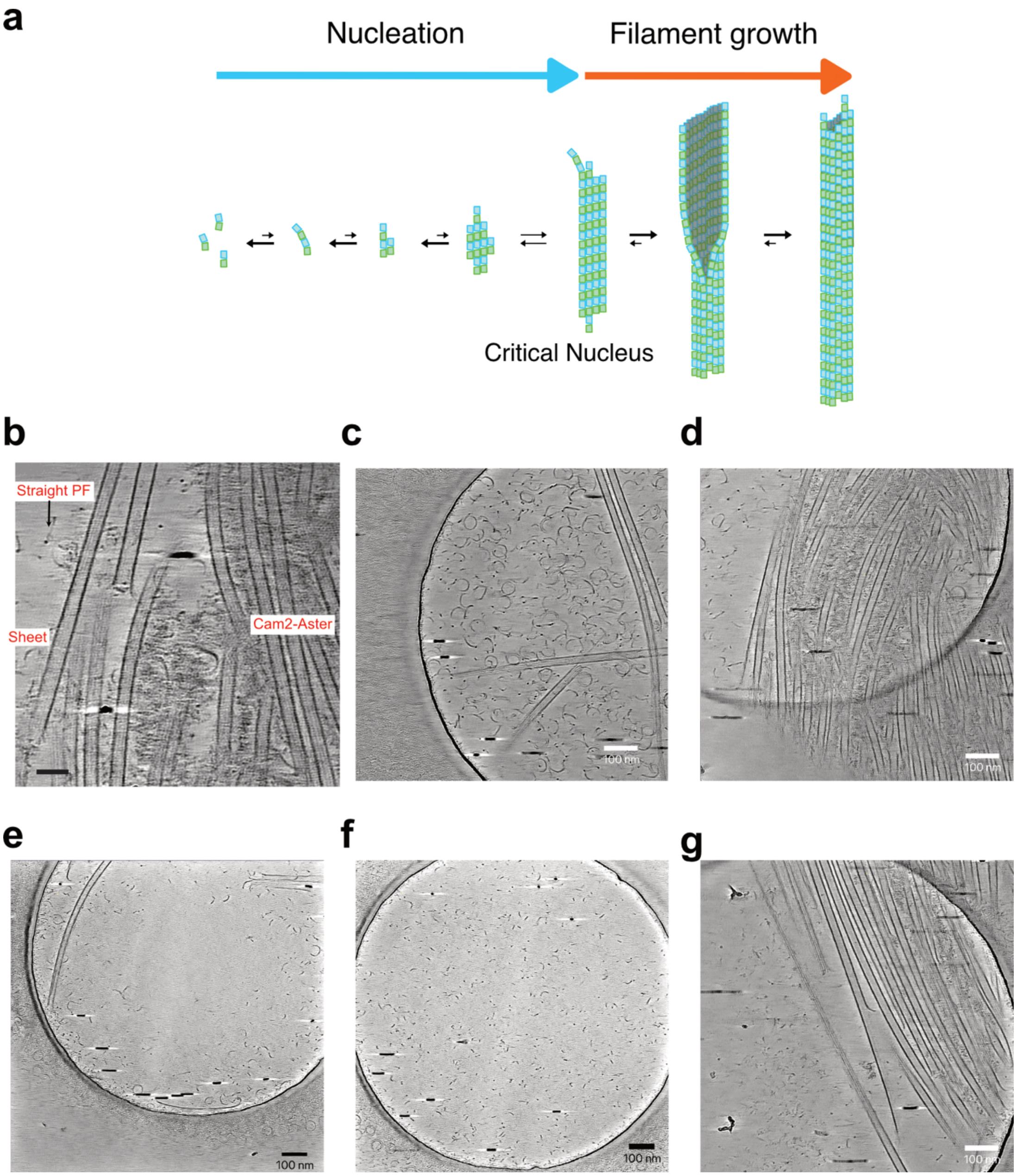

**Supplementary Fig. 1. Cryo-EM visualization of early stages of CAMSAP2-induced microtubule nucleation.** (**a**) Literature-based Model of the spontaneous nucleation during tubulin polymerization into microtubules. (**b**) Cryo-ET reconstruction of 30 μM tubulin with 3 μM CAMSAP2 CC1-CKK after 1 minutes denoised by Cryo-CARE. Microtubules in the Cam2-aster and tubulin sheets and intermediates. Scale bars, 50 nm. (**c–g**) Cryo-electron tomography (cryo-ET) images of CAMSAP2-induced microtubule nucleation after incubation at 37°C for 30 s or 1 min, acquired from different grid holes. (**c**) Tomographic slice corresponding to Fig. 2b (1 min). (**d**) Tomographic slice corresponding to Fig. 2c (1 min). (**e**) Tomographic slice corresponding to Fig. 2D (30 s). (**f**) Representative tomographic slice showing tubulin rings and partial rings (30 s). (**g**) Representative tomographic slice showing an elongating microtubule (1 min).

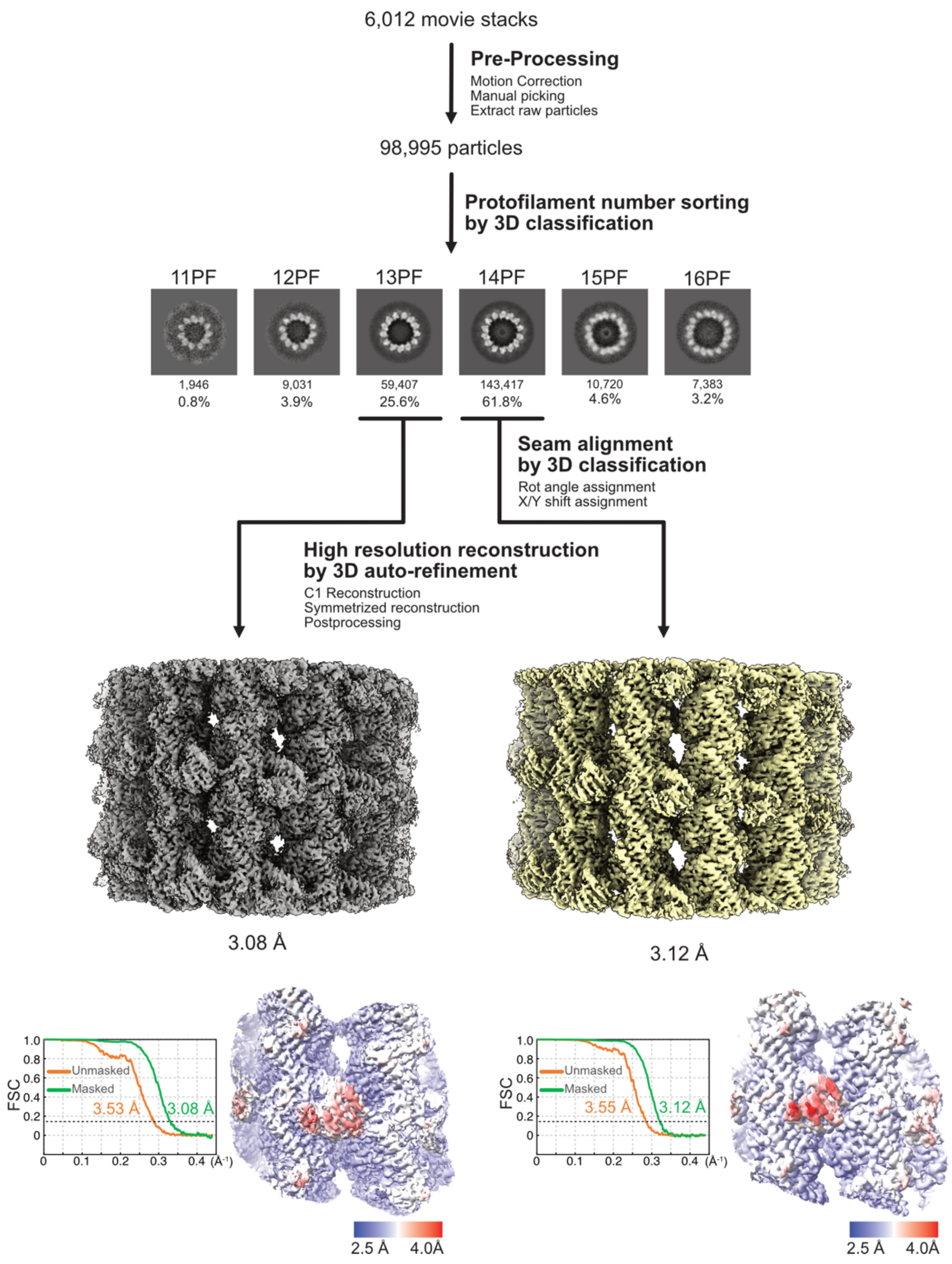

**Supplementary Fig. 2. Outline of the single particle analysis of the CAMSAP2 CC3-CKK.**

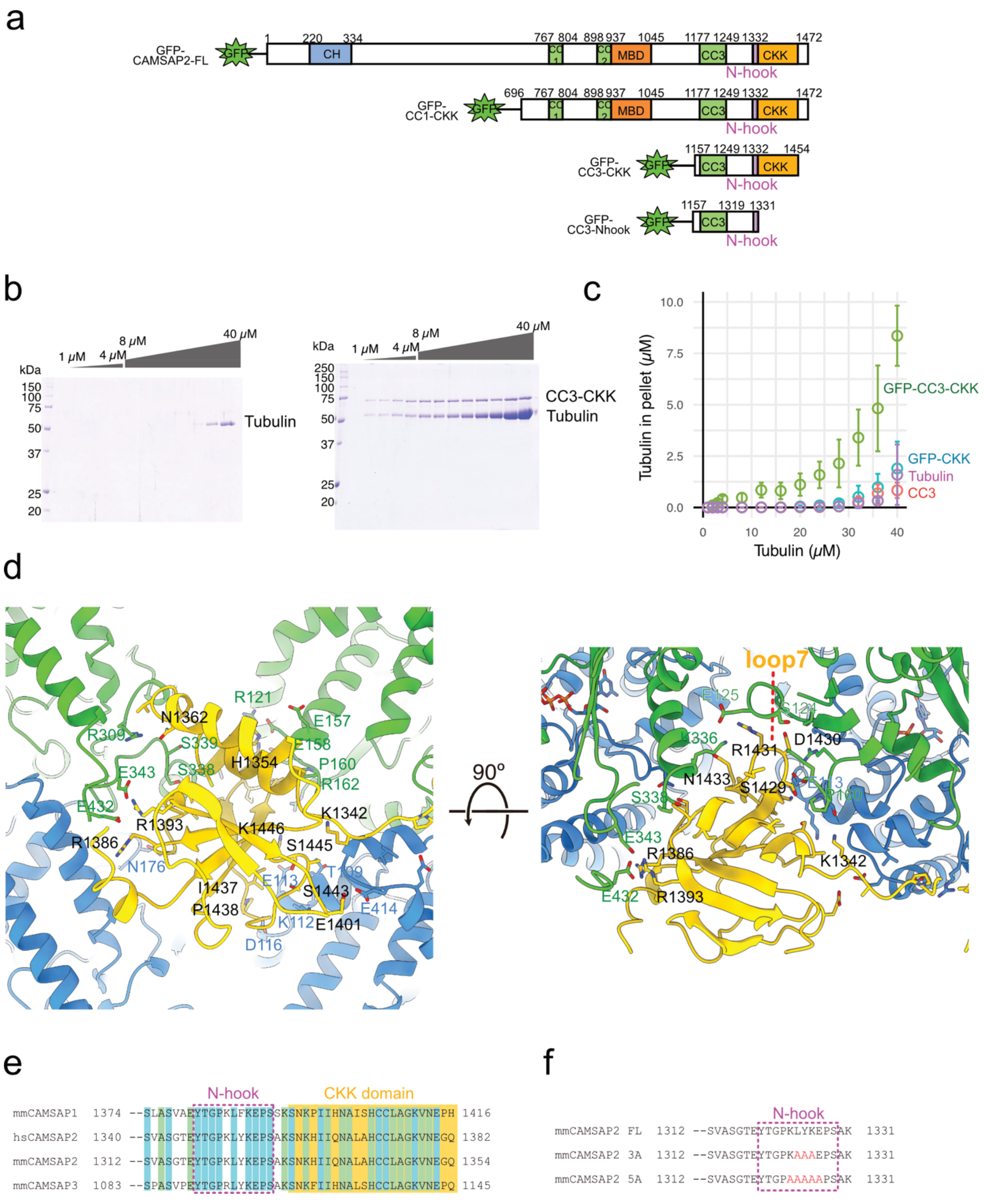

**Supplementary Fig. 3. CAMSAP2 CC3-CKK is a minimum requirement domain for microtubule nucleation.** (**a**) Domain organization of the CAMSAP2 constructs used in this study. (**b**) SDS-PAGE analyses of polymerized tubulin or tubulin with CAMSAP2 CC3-CKK. (**c**) Plots of the depolymerized tubulin concentrations determined by pelleting assay against total tubulin concentration. The average of the three independent experiment assays was plotted. The depolymerized tubulin is purple, CC3 in orange, GFP-CKK in cyan, and GFP-CC3-CKK in light green. (**d**) Molecular interface between CAMSAP2 CKK and microtubule. α-Tubulin is colored green, β-tubulin blue, and CAMSAP2 yellow. (**e**) Sequence alignment between CAMSAPs. (**f**) The design of the series of N-hook mutants. N-hook was mutated to 3 alanines (3A) or 5 alanines (5A).

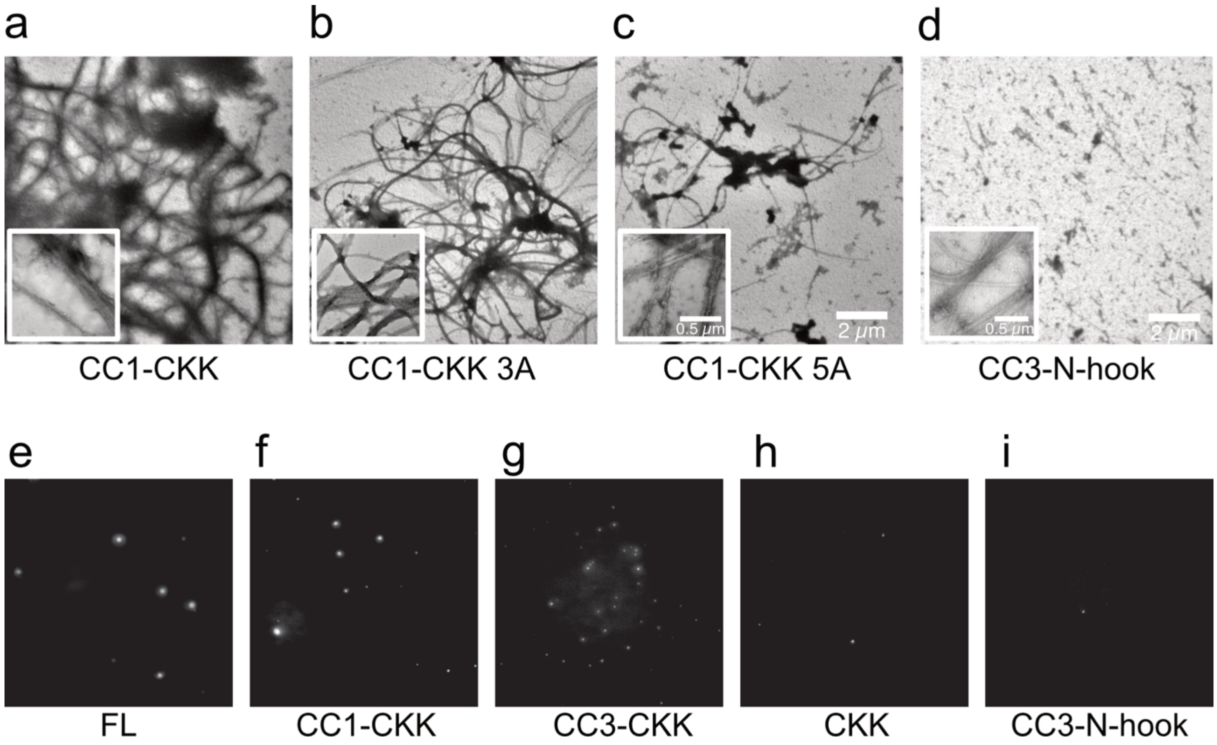

**Supplementary Fig. 4. LLPS and Cam2-aster formation by CAMSAP2 full length and mutants.** (**a**)-(**d**) Cam2-aster formation by CAMSAP2-CC1-CKK, CC1-CKK mutants, and CC3-N-hook. (**e**)-(**i**) LLPS of CAMSAP2 full length and mutants.

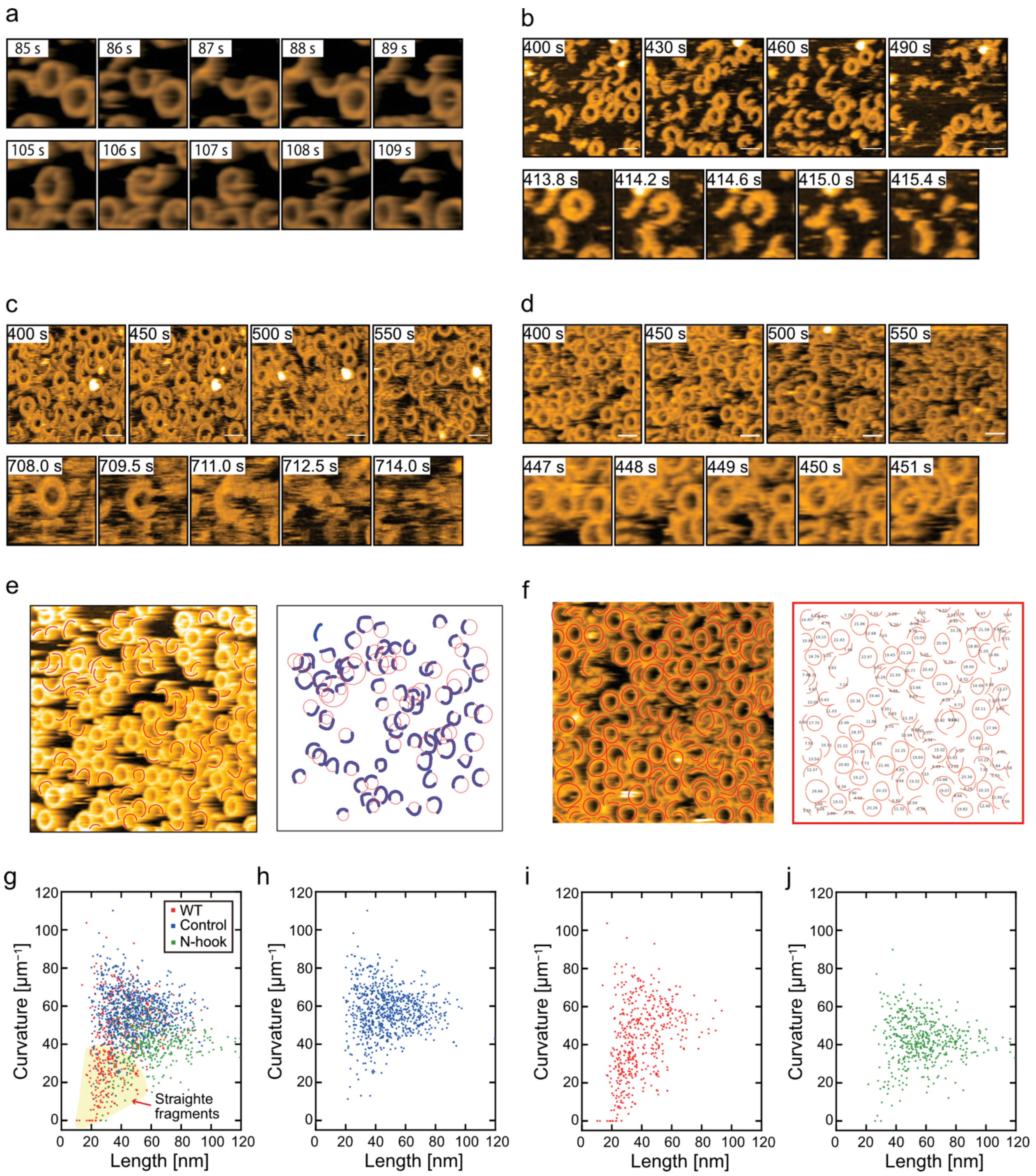

**Supplementary Fig. 5. hsAFM imaging of tubulin rings on mica in the presence and absence of CAMSAP2.** (**a**) Representative hsAFM movie frames showing the protofilaments rings on mica surface. Tubulin rings on the mica surface imaged by hsAFM (see also Supplementary Video 1). (**b–d**) Tubulin rings imaged after addition of (b) CAMSAP2 CC1–CKK 3A mutant, (c) CAMSAP2 CC1–CKK 5A mutant, and (d) CAMSAP2 N-hook, with representative movie frames shown (see also Supplementary Video 2). (**e**) Procedure for estimating total fragment length from AFM images. Left, representative AFM image with tubulin segments manually highlighted in red using Adobe Illustrator. Right, isolated segment representation showing the traced segments with the corresponding interpolated arc lengths indicated in black. Total length was calculated as the sum of all segment lengths. (**f**) Procedure for estimating curvature from AFM images. AFM image of the structure with manually traced contours overlaid. Circular fits obtained using least-squares optimization were applied to each fragment to determine the curvature. (**g-j**) 2D scattering plots showing the distributions of curvature and contour length of tubulin oligomers: control (tubulin only, blue), WT (red), and N-hook (green). The yellow shaded area marks the range defined as “straightened fragments”. Overlayed (e) and individuals (f)-(h).

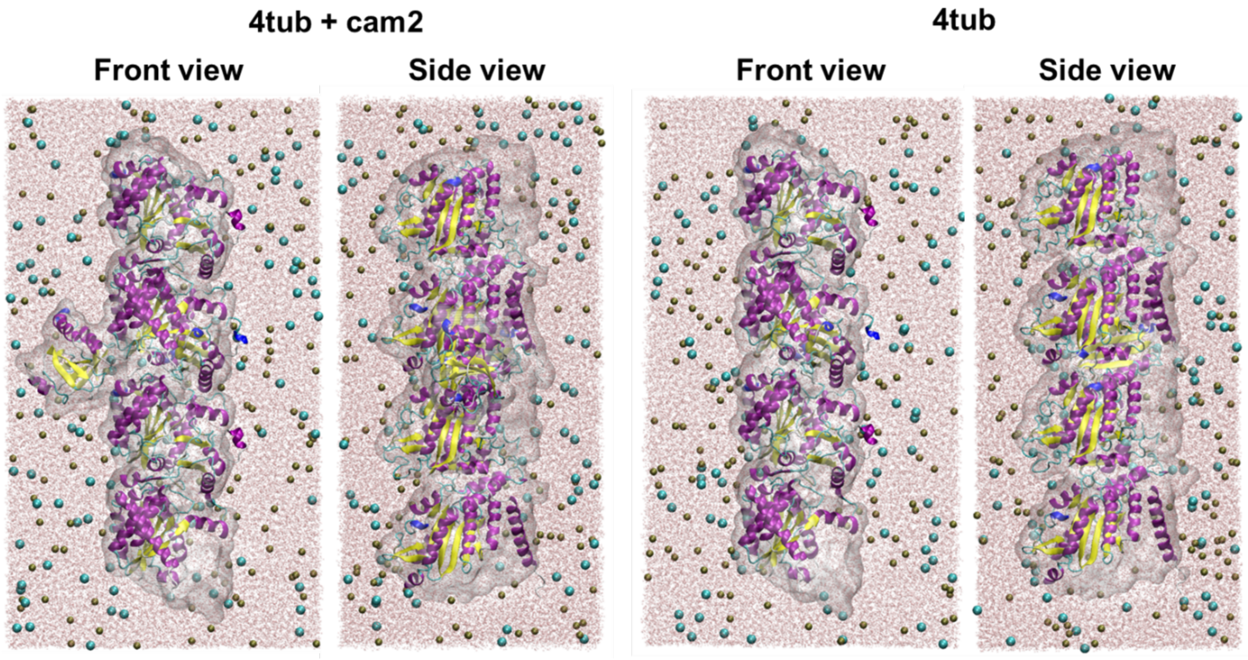

**Supplementary Fig. 6. Visualization for Protein Structure in Explicit-Solvent Simulations.** To highlight the protein structure while retaining solvent context, solvent molecules are selectively displayed in a half-space, revealing the protein interior. The protein is shown as a cartoon, overlaid with a semi-transparent molecular surface on the exposed side, whereas water molecules and ions are rendered as points and spheres, respectively. Panels show orthogonal views of the same configuration.

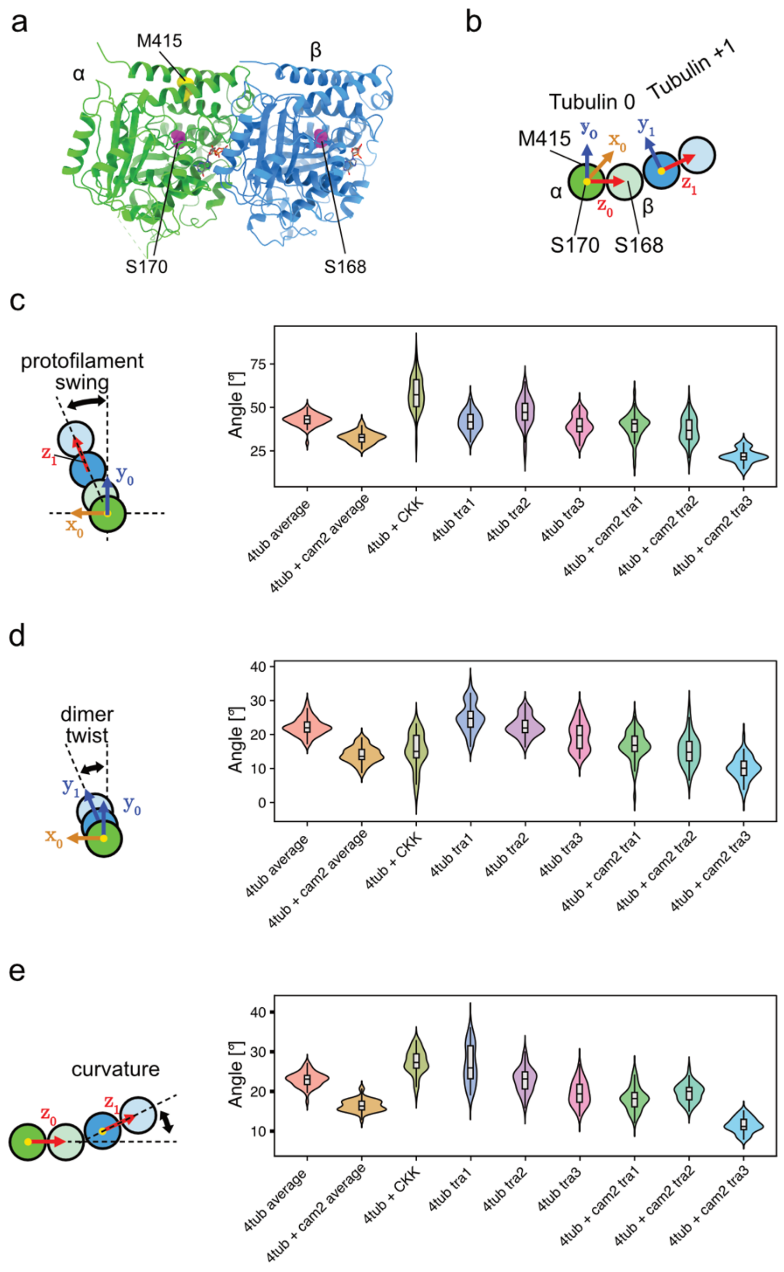

**Supplementary Fig. 7. Distributions of the geometric parameters used to describe tubulin tetramers.** (**a**) Tubulin dimer. Residues used to define the vectors are colored in magenta (α-tubulin S170 and β-tubulin S168) and yellow (β-tubulin M415). (**b**) Vectors defined in the atomic model of a tubulin tetramer. $x_0$ and $x_1$ are calculated as $x_i=y_i \times z_i$, and are therefore perpendicular to the plane of the page. (**c–e**) Violin plots with overlaid box plots showing the distributions of the parameters defined in Fig. 2l, corresponding to those whose mean values are plotted in Fig. 2m-o. Each plot shows the individual values calculated from 50 MD samples per condition: (c) protofilament swing, (d) dimer twist, and (e) curvature.

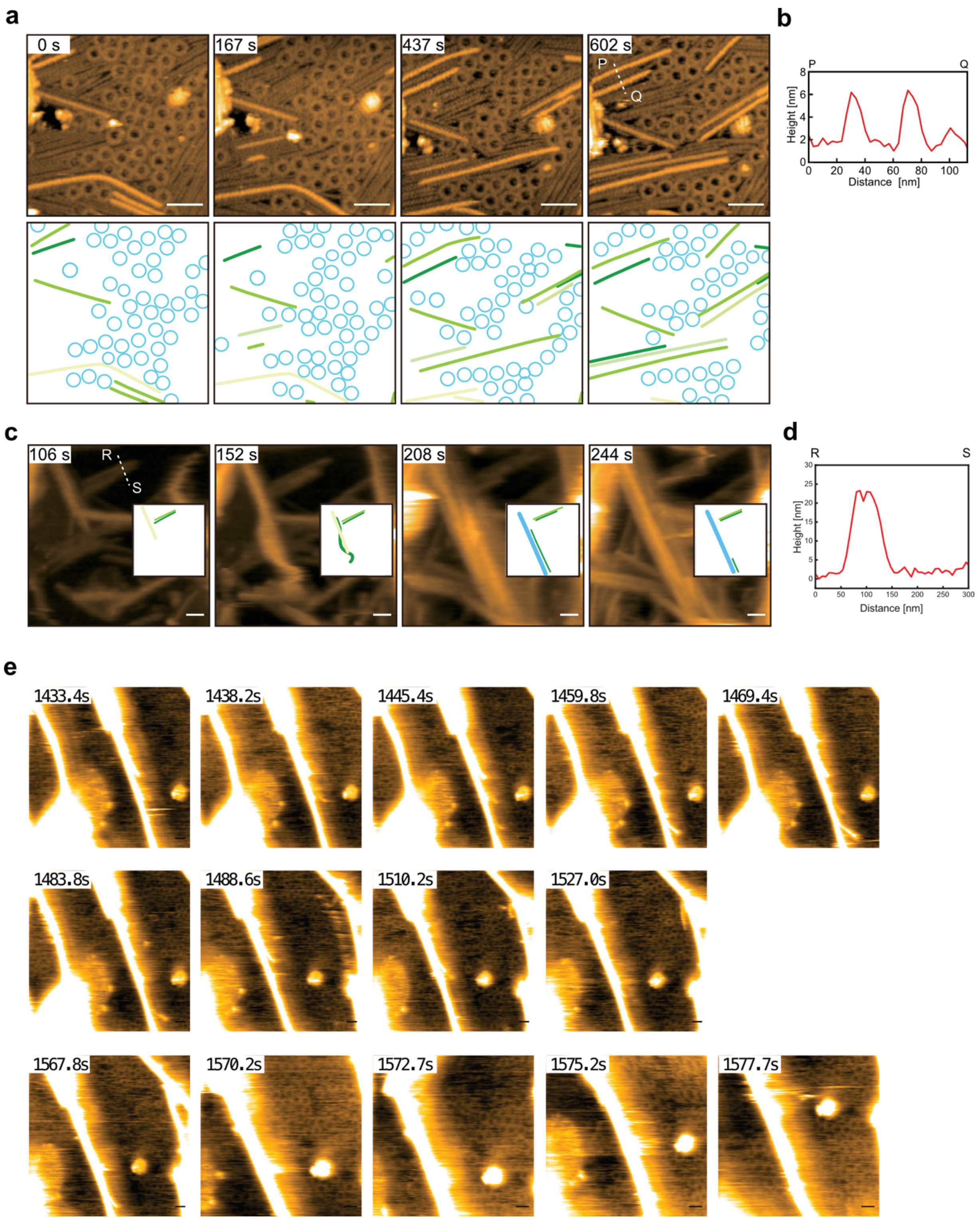

**Supplementary Fig. 8. Representative hsAFM images showing intermediate tubulin structures, protofilament incorporation into sheets, sheet-to-microtubule transitions, and Cam2-aster on a lipid membrane.** (**a**) hsAFM images of tubulin on lipid surface (top) with segmentation (bottom). Protofilaments are color-coded to facilitate tracking across time frames; the same protofilament is shown in the same color in all frames (green, light green, and dark green). Rings are colored in cyan (see also Supplementary Video 3). Protein concentrations of tubulin and CAMSAP2 CC1-CKK were 20 µM and 2 µM. (**b**) Cross-sectional profiles of the heights and widths of the filaments in (a) line P to Q. (**c**) hsAFM images of tubulin on lipid surface with CAMSAP2 CC1-CKK. Dynamic microtubules around a Cam2-aster. Protein concentrations of tubulin and CAMSAP2 CC1-CKK were 20 µM and 2 µM (see also Supplementary Video 3). (**d**) Cross-sectional profiles of the measurable filament heights and widths. (**e**) High contrast image of (c) showing rings in the background. Scale bar, 50 nm.

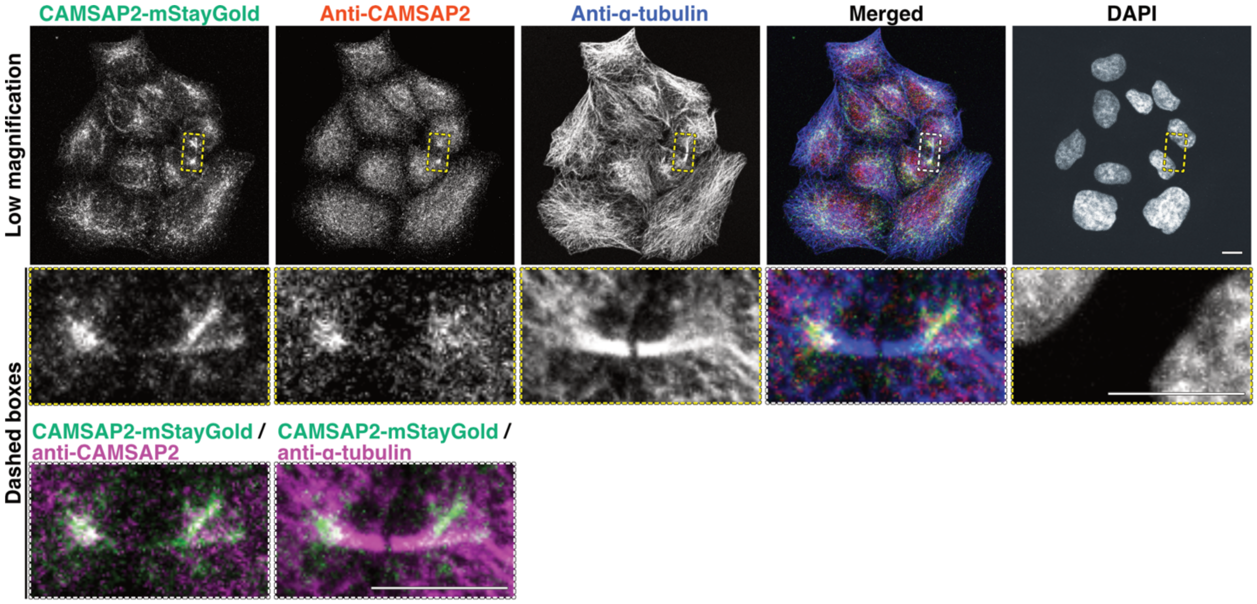

**Supplementary Fig. 9. Endogenous CAMSAP2–mStayGold recapitulates native CAMSAP2 localization in HeLa cells.** Distribution of CAMSAP2–mStayGold (green), endogenous CAMSAP2 (red), and α-tubulin (blue) in mStayGold knock-in HeLa cells. Cells were cultured for 3 days and stained for the indicated proteins; projection images are shown. Dashed boxes indicate regions of interest. Scale bar, 10 µm. The mStayGold tag enabled a clearer and more robust detection of CAMSAP2 than a conventional anti-CAMSAP2 polyclonal antibody.

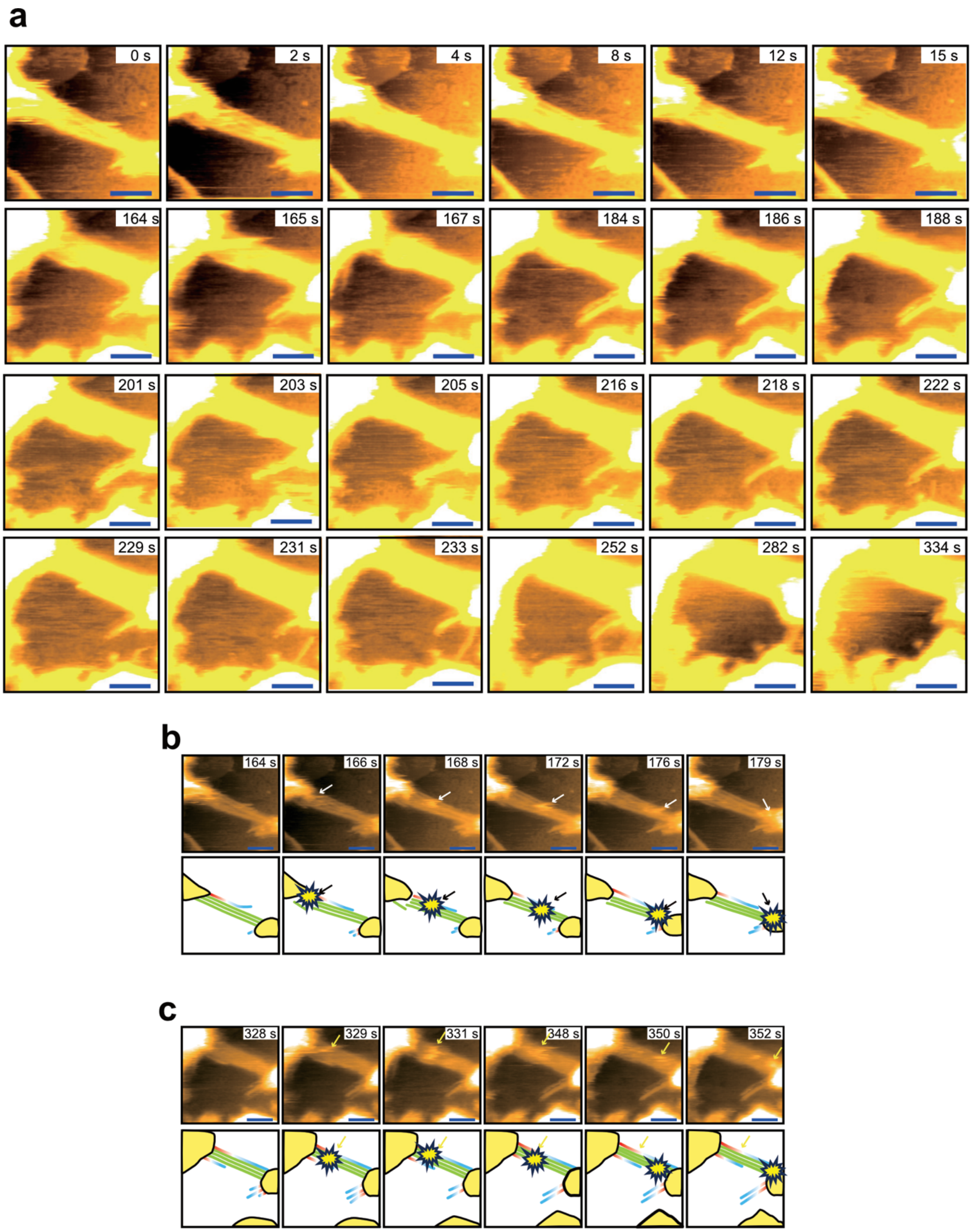

**Supplementary Fig. 10. Cam2-aster network formation observed by hsAFM.** (**a**) High contrast image of Fig. 4b and 4c showing rings and PFs in the background. (**b, c**) Cam2-asters connected by microtubule bundles, with discrete star-shaped oligomeric features moving along the bundles (white and yellow arrows), and corresponding schematics. Microtubule polarity is indicated (minus end, red; plus end, blue). Cam2-asters connected by microtubule bundles, with discrete star-shaped oligomeric features moving along the bundles (white and yellow arrows), and corresponding schematics. Microtubule polarity is indicated (minus end, red; plus end, blue). See also Fig. 4a,b and Supplementary Video 6.

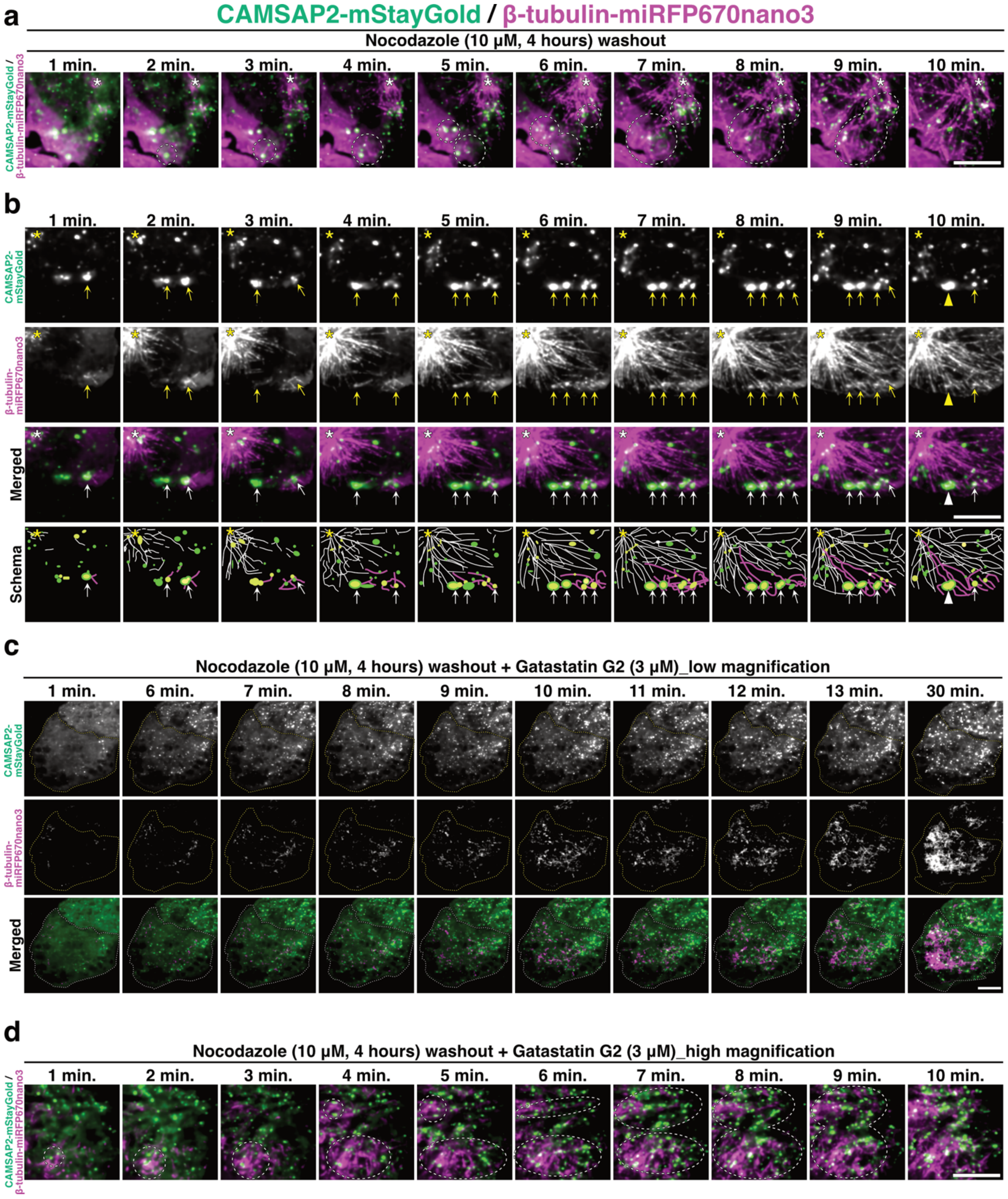

**Supplementary Fig. 11. Microtubular dynamics of the endogenous CAMSAP2, related to Fig. 4.** (**a**) Time-lapse images of microtubule regrowth in HeLa cells carrying endogenous CAMSAP2–mStayGold (green) and β-tubulin–miRFP670nano3 (magenta) knock-ins, shown at 1-min intervals after nocodazole washout. This panel corresponds to a higher-temporal-resolution view of Fig. 4c (right). Cells were treated with 10 μM nocodazole for 4 h and imaged at the basal plane immediately after a 10 min washout. Scale bar, 10 µm. See also Fig. 4c and Supplementary Video 8. (**b**) Representative cell under the same conditions as in (a). Asterisks indicate centrosomes; arrows indicate CAMSAP2 condensates; arrowheads indicate condensate fusion events. White lines denote microtubules elongating from the centrosome, and magenta lines denote microtubules elongating from CAMSAP2 condensates. Scale bar, 10 µm. See also Supplementary Video 9. (**c**) Time-lapse images of microtubule regrowth in a CAMSAP2–mStayGold/β-tubulin–miRFP670nano3 knock-in HeLa cell after co-treatment with 10 µM nocodazole and 3 µM gatastatin G2 for 4 h. Cells were imaged at the basal plane immediately after washout for 30 min at 1-min intervals in the continued presence of 3 µM gatastatin G2. This panel corresponds to higher-temporal-resolution view of Fig. 4b. Scale bar, 10 µm. See also Supplementary Video 10. (**d**) Representative cell under the same conditions as in (c). This panel corresponds to a higher-temporal-resolution view of Fig. 4b (right). Dashed circles indicate regions of interest. Scale bar, 10 µm. See also Fig. 4b and Supplementary Video 11.

#### Table S1. Cryo-EM data collection, refinement and validation statics

| | 13PF MT-CC3-CKK (EMDB-67789) (PDB 21LB) | 14PF MT-CC3-CKK (EMDB-67790) (PDB 21LC) |
|---|---|---|
| **Data collection and processing** | | |
| Magnification | 36,000 | |
| Voltage (kV) | 200 | |
| Electron exposure ($e^{-}$/Å$^2$) | 50 | |
| Defocus range (μm) | −2.5 to −1.0 | |
| Pixel size (Å) | 1.15 | |
| Symmetry imposed | Pseudo-helical | Peudo-helical |
| Initial particle images (no.) | 231,904 | |
| Final particle images (no.) | 59,407 | 143,417 |
| The symmetrized reconstruction<br>Map resolution (Å)<br>FSC threshold | <br>3.08<br>0.143 | <br>3.12<br>0.143 |
| **Refinement** | | |
| Initial model used (PDB code) | 6QUS | |
| Map sharpening *B* factor (Å$^2$) | -62.6 | -92.2 |
| Model composition<br>Non-hydrogen atoms<br>Protein residues<br>Ligands | <br>14,590<br>1,847<br>2 GTP, 2 GDP | <br>14,687<br>1857<br>2 GTP, 2 GDP |
| *B* factors (Å$^2$)<br>Protein<br>Ligand | <br>42.90<br>32.1 | <br>67.46<br>65.22 |
| R.m.s. deviations<br>Bond lengths (Å)<br>Bond angles (°) | <br>0.003<br>0.549 | <br>0.003<br>0.596 |
| Validation<br>MolProbity score<br>Clashscore<br>Poor rotamers (%) | <br>1.80<br>9.58<br>1.78 | <br>1.88<br>11.11<br>1.39 |
| Ramachandran plot<br>Favored (%)<br>Allowed (%)<br>Disallowed (%) | <br>97.44<br>2.56<br>0 | <br>96.69<br>3.31<br>0 |
| Helical parameter<br>Rise (Å)<br>Twist (°) | <br>81.36<br>0.600 | <br>81.38<br>-0.097 |

**Table S2. Latice parameters for 13-pf microtubule structures.**

| SAMPLE | DIMER RISE (Å) | SKEW / TWIST (°) | INTRADIMER RISE (Å) | INTERDIMER RISE (Å) | REFERENCE |
|---|---|---|---|---|---|
| **CAMSAP2-CC3-CKK** | 81.37 | 0.60 | 40.52 | 40.85 | this study |
| **CAMSAP1-CKK-TAXOL** | 82.30 | 0.50 | 41.30 | 41.00 | [26] |
| **GMPCPP** | 83.95 | 0.23 | 41.53 | 42.43 | [55] |
| **GTPΓS** | 81.93 | −0.12 | 41.35 | 40.45 | [55] |
| **GDP** | 81.76 | 0.08 | 41.44 | 40.30 | [55] |
| **GDP-KINESIN** | 81.64 | 0.01 | 41.38 | 40.36 | [55] |
| **GDP-EB3** | 81.44 | −0.12 | 41.40 | 40.30 | [56] |

**Table S3. System sizes and compositions for molecular dynamics simulations.**

| Component | CAMSAP2-bound | CAMSAP2-free | Notes |
|---|---|---|---|
| **α-tubulin** | 2 | 2 | |
| **β-tubulin** | 2 | 2 | |
| **CAMSAP2** | 1 | 0 | bound to central dimer |
| **GTP (α-tubulin)** | 2 | 2 | nucleotide-binding sites |
| **GDP (β-tubulin)** | 2 | 2 | nucleotide-binding sites |
| **$K^+$** | 267 | 275 | 150 mM |
| **$Cl^-$** | 195 | 198 | 150 mM |
| **Water molecules** | 69,066 | 70,028 | TIP3P |
| **Box size ($Å^3$)** | 120×100×210 | 120×100×210 | rectangular box |
| **Total atoms** | 236,861 | 237,630 | |

**Supplementary Video 1**. hsAFM imaging of CAMSAP2 effects on tubulin ring architecture on mica (related to Fig. 2a,c). Left, hsAFM movie showing dynamic changes in tubulin ring curvature in the absence of CAMSAP2 CC1–CKK. Right, hsAFM movie showing tubulin rings in the presence of CAMSAP2 CC1–CKK.

**Supplementary Video 2**. hsAFM imaging of CAMSAP2 mutant effects on tubulin ring architecture on mica (related to Supplementary Fig. 5b-d). hsAFM movies showing dynamic changes in tubulin ring curvature in the presence of CAMSAP2 CC1–CKK 3A (left), CC1–CKK 5A (middle), and N-hook mutant (right).

**Supplementary Video 3**. hsAFM imaging of tubulin ring architecture on a lipid surface without CAMSAP2 (left) or with CAMSAP2 CC1–CKK (right) (related to Supplementary Fig. 8a,c).

**Supplementary Video 4**. hsAFM movies showing slow incorporation of protofilaments into tubulin sheets via inchworm-like straightening and winding (related to Fig. 3a).

**Supplementary Video 5**. hsAFM movies revealing stepwise polymerization assisted by the CAMSAP2 CC1–CKK 5A mutant (related to Fig. 3d).

**Supplementary Video 6**. hsAFM movies of Cam2-aster showing dynamic behavior of asters and microtubules (related to Fig. 4a,b, Supplementary Fig. 10).

**Supplementary Video 7**. Microtubule regrowth dynamics in HeLa cells with endogenous CAMSAP2–mStayGold (green) and β-tubulin–miRFP670nano3 (magenta) double knock-in (related to Fig. 4c left). Cells were treated with 10 μM nocodazole for 4 h and imaged at the basal plane immediately after washout for 30 min at 1-min intervals. Scale bar, 10 μm.

**Supplementary Video 8**. Higher-magnification view of microtubule regrowth dynamics in HeLa cells with endogenous CAMSAP2–mStayGold (green) and β-tubulin–miRFP670nano3 (magenta) double knock-in (related to Fig. 4c right and Supplementary Fig. 11a). Cells were treated with 10 μM nocodazole for 4 h and imaged at the basal plane immediately after washout for 10 min at 1-min intervals. Scale bar, 10 μm.

**Supplementary Video 9**. CAMSAP2 condensation and associated microtubule network formation in a HeLa cell with endogenous CAMSAP2–mStayGold (green) and β-tubulin–miRFP670nano3 (magenta) double knock-in (related to Supplementary Fig. 11b). Cells were treated with 10 μM nocodazole for 4 h and imaged at the basal plane immediately after washout for 10 min at 1-min intervals. Scale bar, 10 μm.

**Supplementary Video 10**. Microtubule regrowth dynamics in HeLa cells with endogenous CAMSAP2–mStayGold (green) and β-tubulin–miRFP670nano3 (magenta) double knock-in following co-treatment with 10 μM nocodazole and 3 μM Gatastatin G2 for 4 h (related to Fig. 4d left and Supplementary Fig. 11c). Cells were imaged at the basal plane immediately after washout for 30 min at 1-min intervals in the continued presence of 3 μM Gatastatin G2. Scale bar, 10 μm.

**Supplementary Video 11**. Higher-magnification view of microtubule regrowth dynamics in a HeLa double knock-in cell expressing endogenous CAMSAP2–mStayGold (green) and β-tubulin–miRFP670nano3 (magenta) following co-treatment with 10 μM nocodazole and 3 μM Gatastatin G2 for 4 h (related to Fig. 4d right and Supplementary Fig. 11d). Cells were imaged at the basal plane immediately after washout for 10 min at 1-min intervals in the continued presence of 3 μM Gatastatin G2. Scale bar, 10 μm.
